# Multiplatform Modeling of Atrial Fibrillation Identifies Phospholamban as Central Regulator of Cardiac Rhythm

**DOI:** 10.1101/2022.09.23.509238

**Authors:** Anaïs Kervadec, James Kezos, Haibo Ni, Michael Yu, Sean Spiering, Suraj Kannan, Peter Andersen, Eleonora Grandi, Karen Ocorr, Alexandre R. Colas

## Abstract

Atrial fibrillation (AF) is the most common form of sustained cardiac arrhythmia in humans, present in > 33 million people worldwide. Although AF is often developed secondary to cardiovascular diseases, endocrine disorders, or lifestyle factors, recent GWAS studies have identified >200 genetic variants that substantially contribute to AF risk. However, it is currently not known how these genetic predispositions contribute to the initiation and/or maintenance of AF-associated phenotypes. In this context, one major barrier to progress is the lack of experimental systems enabling to rapidly explore the function of large cohort of genes on rhythm parameters in models with human atrial relevance. To address these modeling challenges, we have developed a new multi-model platform enabling *1)* high-throughput characterization of the role of AF-associated genes on action potential duration and rhythm parameters at the cellular level, using human iPSC-derived atrial-like cardiomyocytes (ACMs), and at the whole organ level, using the Drosophila heart model, and *2)* validation of the physiological relevance of our experimental results using computational models of heterogenous human adult atrial myocytes (HAMs) and tissue. As proof of concept, we screened a cohort of 20 AF-associated genes and identified Phospholamban (PLN) loss of function as a top conserved hit that significantly shortens action potential duration in ACMs, HAMs and fly cardiomyocytes. Remarkably, while PLN knock-down (KD) was not sufficient to induce arrhythmia phenotypes, addition of environmental stressors (*i.e* fibroblasts, β-adrenergic stimulation) to the model systems, led to the robust generation of irregular beat to beat intervals, delayed after depolarizations, and triggered action potentials, as compared to controls. Finally, to delineate the mechanism underlying PLN KD-dependent arrhythmia, we used a logistic regression approach in HAM populations, and predicted that PLN functionally interacts with both NCX (loss of function) and L-type calcium channels (gain of function) to mediate these arrhythmic phenotypes. Consistent with our predictions, co-KD of PLN and NCX in ACMs and flies, led to increased arrhythmic events, while treatment of ACMs with L-type calcium channel inhibitor, verapamil, reverted these phenotypes. In summary, these results collectively demonstrate that our integrated multi-model system approach was successful in identifying and characterizing conserved roles (*i.e* regulation of Ca2+ homeostasis) for AF-associated genes and phenotypes, and thus paves the way for the discovery and molecular delineation of new gene regulatory networks controlling atrial rhythm with application to AF.

## INTRODUCTION

Atrial fibrillation (AF) is the most common form of sustained cardiac arrhythmia in humans (Du et al., 2017). At the whole heart level, a central feature of AF is a very rapid and uncoordinated atrial activity while at the cellular level, the mechanism maintaining arrhythmia often arises from a “vulnerable substrate”, which consists of either action potential duration (APD) prolongation or shortening events (Christophersen et al., 2013). Such vulnerable substrates are thought to be caused by genetic predispositions, cardiac remodeling caused by heart disease, aging, and/or altered regulation by neurohormonal factors (Campuzano and Brugada, 2009; Chen et al., 2014; Lip et al., 2016; Weng et al., 2017). In this context, linkage analysis in familial cases of AF(Chen et al., 2003; Hodgson-Zingman et al., 2008; Olson et al., 2006) as well as genome-wide associated studies (GWAS) in the general population (Christophersen et al., 2017; Nielsen et al., 2018b; Roselli et al., 2018), have elucidated some of the genetic underpinnings associated with the disease. As a result, close to 140 genetic loci linked to >200 genes have been identified (Fatkin et al., 2017; Roselli et al., 2018; Roselli et al., 2020), however none of these genes have been validated as disease-causing in the general population, limiting drug discovery efforts. In this context, a major barrier to progress is the lack of experimental platforms/strategies enabling rapidly establishment of causal links between gene function and AF-associated phenotypes (electrical remodeling, arrhythmia). Among the variety of models available to evaluate AF-associated genes, the four-chambered mouse heart has been extensively used to establish functional links between genes or genetic loci and rhythm phenotypes (Lozano-Velasco et al., 2016; Nadadur et al., 2016; Temple et al., 2005; van Ouwerkerk et al., 2019; Wang et al., 2010; Zhang et al., 2019). However, despite high proteome homology with humans and ability to manipulate the genome, the substantial electrophysiological differences (fast resting rate, short AP duration and triangular shape, species-specific K+ channels (Kaese and Verheule, 2012)), relatively long lifespan (years) and low throughput capacity of methods to retrieve electrophysiological parameters, limit the use of mice as a primary model for gene discovery related to AF.

In contrast to mice, flies have a short generation time (∼10 days) and established automated kinetic imaging techniques (Fink et al., 2009; Klassen et al., 2017), coupled with available functional genomic resources (e.g. Flybase.org; VDRC(Mohr et al., 2014)), enable the rapid evaluation of gene function on rhythm parameters at the whole heart level. In addition, although the fly heart structurally differs from that in vertebrates, the fundamental mechanisms of development and function are remarkably conserved, including a common transcriptional regulatory network (Bodmer, 1995; Cripps and Olson, 2002), shared protein composition (Cammarato et al., 2011), and electrical and metabolic properties (Diop and Bodmer, 2015; Ocorr et al., 2007b; Ocorr et al., 2014). Thus, the adult fly heart represents an appealing model to rapidly evaluate the role of evolutionarily conserved genes for their ability to regulate cardiac rhythm, although a limitation to this model is the lack of atrial specificity.

The advent of iPSC technology (Takahashi et al., 2007; Takahashi and Yamanaka, 2006) and protocols enabling the generation of subtype-specific cardiomyocytes (Burridge et al., 2014; Cunningham et al., 2017; Devalla et al., 2015; Yu et al., 2018), provide a unique experimental access to human atrial myocyte biology. In addition, the recent development of high-throughput (HT) kinetic imaging techniques (Cerignoli et al., 2012; McKeithan et al., 2017), fluorescent calcium and voltage-sensing indicators (Liu and Miller, 2020; Paredes et al., 2008), coupled with available functional genomic resources (siRNAs, CRISPR/Cas9 guides libraries), enable large-scale exploration of gene function on human cardiac electrophysiological (voltage and calcium transients kinetics) and rhythm (rhythmicity, beat rate) parameters (Elmen et al., 2020; Murphy et al., 2021). Although human iPSC-derived atrial-like cardiomyocytes (ACMs) are well suited to identify atrial-specific and cell autonomous rhythm-regulating mechanisms (Devalla et al., 2016; Marczenke et al., 2017a; Marczenke et al., 2017b), the relative immaturity of hiPSCs-derived CMs (Cho et al., 2017; Yang et al., 2014) and inherent lack of tissue level integration, might limit translation of the findings to the adult human heart. In sum, single model approaches are limited in their ability to validate large cohorts of AF-associated genes, indicating the necessity to develop alternative strategies to improve AF gene validation.

Based on these observations, we reasoned that combining assays with human, atrial and whole organ relevance that also have HT functional genomics capacity could enhance our ability to rapidly establish causal links between AF-associated genes and arrhythmia phenotypes. To establish such a platform, we developed a human-relevant assay that measures APD in ACMs with single cell resolution. In parallel, we optimized a fly cardiac function assay that measures contraction duration (systolic interval (SI)), as a surrogate measurement for APD. As proof of concept, we screened a cohort of 20 AF-associated genes and identified Phospholamban (PLN) loss of function as a top conserved hit that significantly shortens action potential duration in ACMs, HAMs and fly cardiomyocytes. Remarkably, while PLN knock-down (KD) was not sufficient to induce arrhythmia phenotypes, addition of environmental stressors (i.e fibroblasts, β-adrenergic stimulation) to the model systems, led to the robust generation of irregular beat to beat intervals, delayed after depolarizations, and triggered action potentials, as compared to controls. Finally, to delineate the mechanism underlying PLN KD-dependent arrhythmia, we used a logistic regression approach in HAM populations, and predicted that PLN functionally interacts with both NCX (loss of function) and L-type calcium channels (gain of function) to mediate these arrhythmic phenotypes. Consistent with our predictions, co-KD of PLN and NCX in ACMs and flies, led to increased arrhythmic events, while treatment of ACMs with L-type calcium channel inhibitor, verapamil, reverted these phenotypes.

## RESULTS

### Integrated Multi-Model Systems Platform to Identify Rhythm-Regulating Genes

To phenotypically assess AF-associated genes, we established a novel phenotypic platform enabling to study gene function on APD and rhythm parameters in HT in both ACMs and flies.

### Single cell and HT assessment of APD and rhythm parameters in ACMs

To study the molecular basis of chamber-specific electrical disorders such as atrial fibrillation, Id1-overexpressing cardiac progenitors (CPs) were used to generate ACMs as described in (Cunningham et al., 2017; Yu et al., 2018). Treatment of Id1-induced CPs with a single dose of retinoic acid (300 nM) efficiently promoted the generation of atrial-like, NR2F2+ beating CMs (∼ 80% were NR2F2+, ACTN2+) (**Fig. 1A-C**). Consistent with an atrial-like identity, induced ACMs also expressed atria-enriched (Devalla et al., 2015; Uhlen et al., 2015) transcription factors (NR2F2, TBX5, ZNF385B), ion channel genes KCNA5 (encoding Kv1.5) and KCNJ3 (encoding Kir 3.1), ligands (NPPA, NPPB) and receptors (EGR1/2, PDGFRA), at both day 12 and at day 25 of differentiation (**Fig. 1D and Suppl. Fig. 1A,B**). Electrophysiologically, ACMs typically generated short (∼120 ms) and triangular action potentials (**Fig. 1E**) while untreated Id1-CPs generated CMs displayed longer action potential (∼200 ms) with a plateaued phase 2 (**Suppl. Fig. 1C**), reminiscent of a ventricular-like identity (Ng et al., 2010). Moreover, and consistent with an atrial cell fate, ACMs also displayed shorter calcium transient durations (CTD_50_ and CTD_75_) as compared to ventricular CMs (VCMs) (**Suppl. Fig. 1D-G**) (Ng et al 2010).

**Figure 1.**
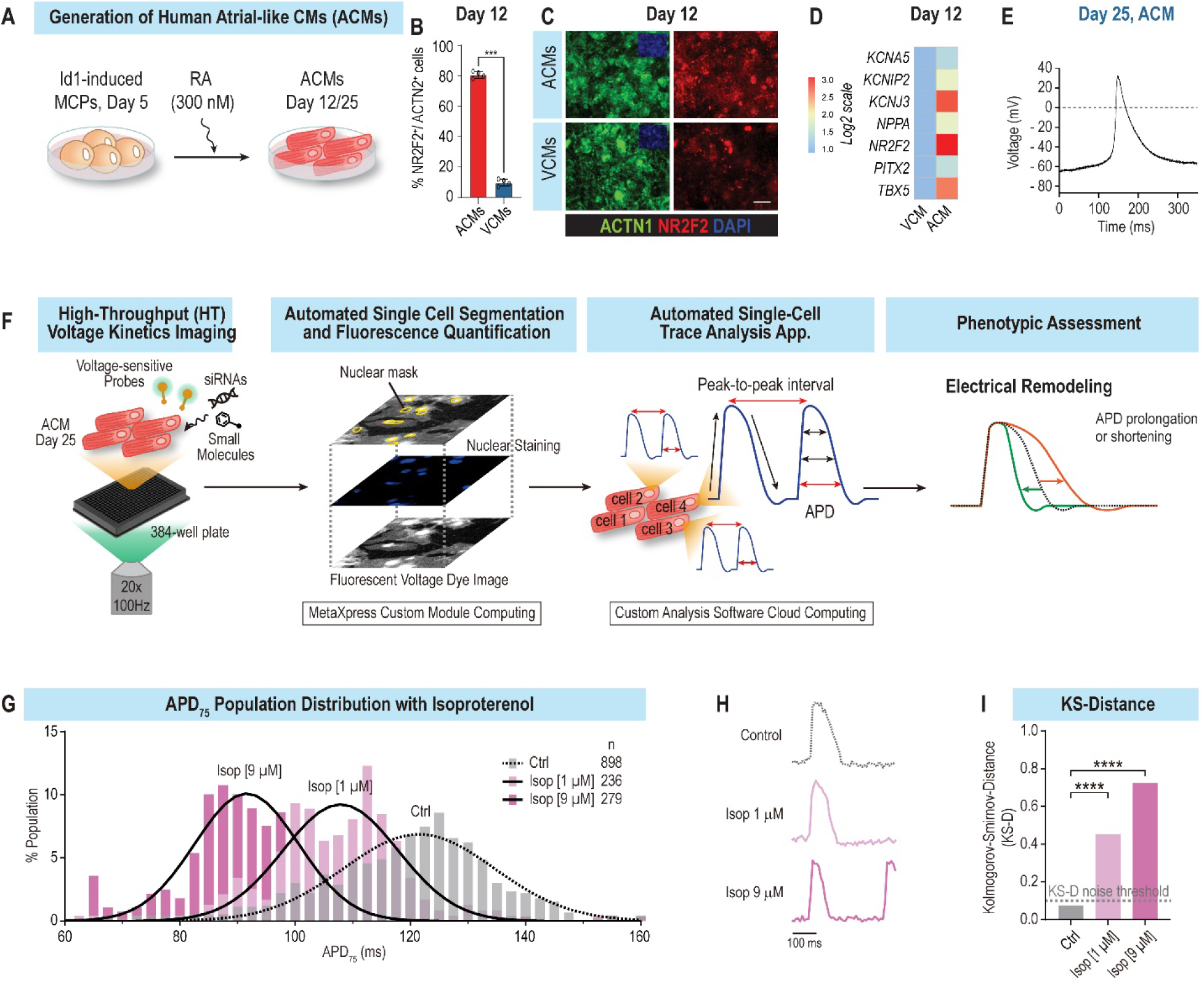
ACM platform. **(A)** Schematic representation of the ACM differentiation protocol. To promote atrial differentiation, day 5 cardiac progenitors were treated with 300 nM retinoic acid and subsequently cultured until either at day 12 or 25. **(B)** RA treatment efficiently induces the generation of atrial-like NR2F2+ beating CMs (∼80% of NR2F2+, ACTN1+). **(C)** Representative immunofluorescence images showing overexpression of NR2F2+ (red) and ACTN1+ (Green) cells in ACMs **(D)** Heatmap of atrial genes enriched in day 12 ACMs as compared to VCMs. **(E)** Patch-clamp experiments show that ACMs generate atrial-like triangular-shaped action potentials. **(F)** Schematic representation of single cell and HT platform to measure APD parameters in ACMs. **(G)** Population distribution of APD_75_ values from ACMs treated with escalating doses of isoproterenol, show a dose dependent APD shortening. **(H)** Single AP traces of Median APD_75_ for each condition. **(I)** Histogram representing Kolmogorov-Smirnov Distance values (KS-D) for Ctrl and isoproterenol treated ACMs.

Next, to facilitate the characterization of AF-associated arrhythmia phenotypes in ACMs, we developed an imaging platform that automatically tracks and quantifies action potential (AP) and rhythm parameters in HT with single cell resolution (**Fig. 1F**). To retrieve AP and rhythm parameters, ACMs were co-labeled with a voltage dye (VF2.1Cl) and a nuclear dye (Hoechst 33258) as described in (McKeithan et al., 2017). For each condition, one image of the Hoechst dye was collected followed by a 5-seconds acquisition of the voltage dye channel at 100 Hz. Next, using a custom algorithm developed on the ImageXpress, each cell in the field of view was segmented using the Hoechst topological information and each cell mask was propagated to the “voltage dye channel”, thereby enabling the quantification of voltage-dependent fluorescence variation over time with single-cell resolution. To retrieve the electrophysiological parameters, we developed a cloud-based trace analysis application that automatically processes each AP trace and retrieves median and standard deviation values, for APD-10, 25, 50, 75, 90; T25-75, T75-25; Vmax up and down; beat rate; peak-to-peak interval; and rhythm regularity index; for each cell. This platform enabled us to automatically record, quantify and analyze AP and rhythm parameters in less than 2 minutes per condition.

To test the platform’s ability to identify APD modulators, we infused ACMs with isoproterenol, a non-selective β-adrenergic agonist, known to both shorten APD and increase beat rate in hiPSC-CMs (McKeithan et al., 2017). Consistent with previous studies, escalating doses of isoproterenol caused a dose-dependent shortening of median APD_75_ values, from 121.3 ms (untreated) to 108.6 ms (1 µM) and 91.2 ms (9 µM) (**Fig. 1G,H**). To measure whether the isoproterenol treatment has a significant effect on APD at the whole cell population level, we used the Kolmogorov-Smirnov test (KS-D) (Dal Molin et al., 2017; Delmans and Hemberg, 2016; Feng et al., 2009; Gaber et al., 2013), which enabled us to quantify and compare the distributional differences of binary features such as APD_75_ or beat rate. Consistent with median APD_75_ values, escalating doses of isoproterenol (1 and 9 µM) led to an increase of APD_75_ KS-D value to ∼0,4 and 0.7 respectively, as compared to control (**Fig. 1I**). Similarly, to quantify cellular manifestations of arrhythmia, we defined an arrhythmia index (AI) that quantifies beat-to-beat interval irregularities as a metric for arrhythmically beating cells (**Suppl. Fig. 3A**). In this context, we assess that AP trains with AI values lower than 20, generally describe regular beating patterns, while AP train with AI values >20 mark generally marks irregular beating patterns (**Suppl. Fig. 3B**). Finally, to benchmark our platform for arrythmia-associated phenotypes, we tested the role of dofetilide, a class III anti-arrhythmic, that selectively blocks the rapid component of the delayed rectifier outward potassium. At therapeutic dose, dofetilide prolongs APD and subsequently increase of the refractory period, thereby mediating its anti-arrhythmic effect (Geng et al., 2020). Conversely, at higher doses dofetilide increases the incidence of arrhythmia phenotypes such as early afterdepolarizations (EADs) (Jaiswal and Goldbarg, 2014; McKeithan et al., 2017). Consistent with these observations, ACMs treated with a low dose of dofetilide (33 nM) displayed a reduced AI as compared to control, while high dose of dofetilide (100nM) dramatically increased the percentage of cells with AI values > 20 (from 4% to 67%) and associated KS-D value of 0.7 (**Suppl. Fig. 3C-F**). Collectively, these data show that this new phenotypic platform enables the HT and automated quantification of APD and rhythm parameters in ACMs with single cell resolution.

### HT measurement of APD and rhythm in Drosophila cardiac platform

To assess AF-associated mechanisms at the whole organ level using the Drosophila model, we used high-speed video recording of heart movements in *in situ* preparations. Heart function was quantified as previously described (Fink et al., 2009; Vogler and Ocorr, 2009); http://sohasoftware.com/) providing precise measurements of heart period (R-R interval), systolic interval (SI), as well as arrhythmicity and fractional shortening/contractility in a functioning heart. We have previously shown that most of the key cardiac ion channels present in human hearts are also present and functional in the fly heart (Ocorr et al., 2007b; Ocorr et al., 2017) and **Supplemental Table 1**). Importantly, our previous studies using simultaneous optical and intracellular recordings demonstrated a direct 1 :1 correlation between myocardial cell depolarization and heart wall movement. It is important to note that the fly heart is composed of a single layer of myocardial cells and any heart wall movement is an immediate reflection of the contractile state of component myocardial cells. Thus, we quantified APD and the corresponding systolic interval (SI) from simultaneous electrical and optical recordings respectively from hearts of middle-aged wildtype controls and KCNQ mutants and found good agreement between APD and SI (**Fig. 2 B-E**). Therefore, we used cardiac contraction / relaxation movements as surrogates for APD (Cammarato et al., 2015). Cardiac-specific gene KD was achieved using a Gal-4 based system (Brand and Perrimon, 1993) that drives expression of dsRNAi in a cardiac-specific manner. Since AF is an aging related disease, the fly provides an opportunity to examine effects of cardiac gene KD at young, middle and old ages (∼1, 3 and 5 weeks respectively).

**Figure 2.**
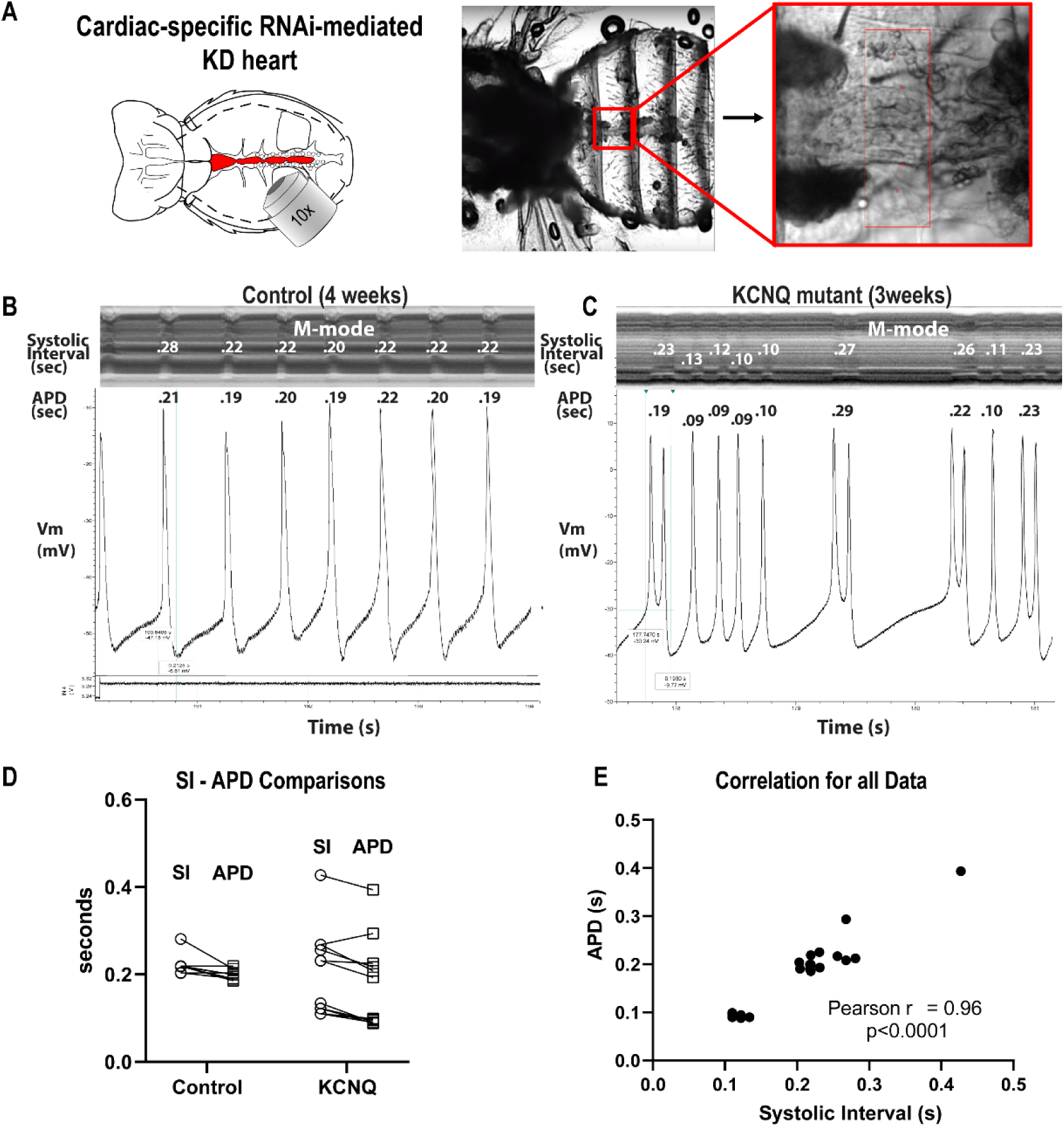
Fly heart platform. **(A)** Schematic of the fly thorax and abdomen (left, heart tube is shown in red.) and image of semi-intact preparation (middle) with a single cardiac chamber (red box) shown at higher magnification to the right. **(B & C)** Simultaneous optical and electrophysiological recordings from beating hearts. M-modes from optical recordings are shown on the top with the corresponding Action Potential (AP) traces below. AP duration (APD) and Systolic Intervals are shown in seconds. The lower window in B shows the voltage trace generated by the image capture software that was used to synchronize the optical and electrical recordings. **(D)** Systolic intervals are paired with their corresponding APs. **(E)** The Pearson Correlation Coefficient for the combined data in D showed a significant correlation between SIs and APDs (r = 0.96, p<0.0001).

### Functional screen of AF-associated genes identifies PLN loss of function as major driver of APD and contraction intervals shortening

To evaluate the ability of the platform to identify AF-associated genes and mechanisms, we first assessed the expression of genes previously associated with AF (Fatkin et al., 2017), by RNA-seq of day 12 and day 25 ACMs. The result revealed that most AF candidate genes were expressed in ACMs at moderate-to-high levels (from 0.1 to >100 RPKM) (**Suppl Fig. 3A**) and most are also expressed in the fly heart (Supplemental Table 1). Next, we selected 20 genes that had been identified in rare variant familial AF studies and/or as having SNPs reported in GWAS studies (**Suppl Fig. 3B and (Fatkin et al., 2017)**). To evaluate their effect on APD, we transfected siRNAs directed against these 20 AF-associated genes into day 25 ACMs and measured voltage variation over time with single cell resolution. Remarkably, APD_75_ population measurements revealed that 9 out of 20 siRNAs, robustly induced electrical remodeling (KS-D value > 0.25, p-value < 0.001) (**Fig. 3A and Suppl Fig. 3C**). Interestingly, the screen identified two phenotypes: prolongation or shortening of APD. Among these, down-regulation of GATA5, GATA6, PITX2, and KCNA5 significantly prolonged APD, whereas KD of PLN and KCND3 shortened APD. Notably, none of the siRNA knockdowns alone were able to induce overt arrhythmia-like phenotype (*i.e.* a significant percentage of cells with AI > 20) in ACMs (data not shown).

**Figure 3.**
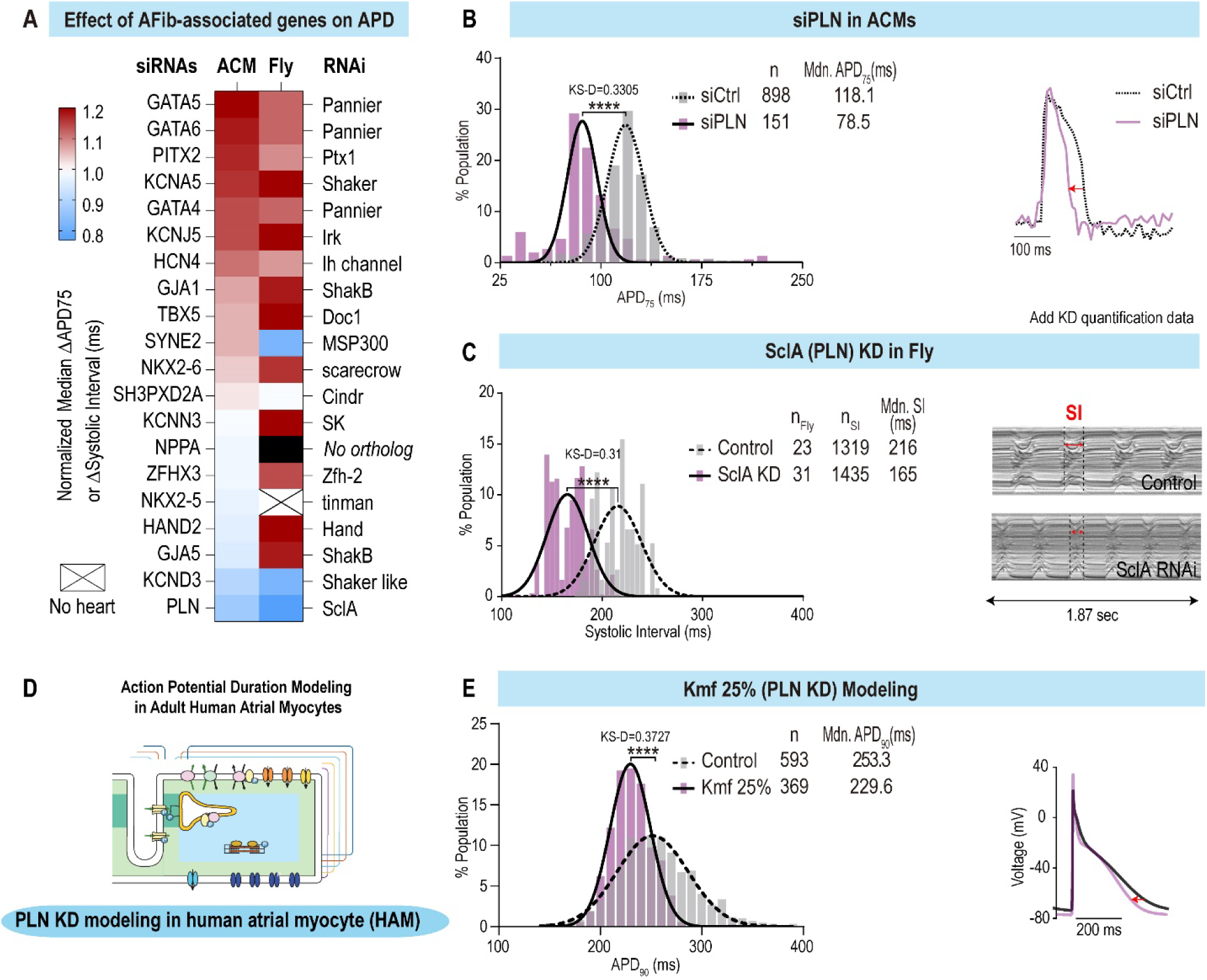
Loss of function screen of AF-associated genes identifies conserved modulators of APD and SI in multi-model system platform. **(A)** Heatmap showing the normalized effects of AF-associated genes loss of function on APD and SI in ACMs and flies respectively. **(B)** Population distribution of APD_75_ values for siControl and siPLN transfected ACMs (left) and representative AP traces (right) showing the shortening effect of siPLN. **(C)** Population distribution of SIs in control vs SclA KD conditions in flies (left) and representative m-modes (right) showing the SI shortening effect for SclA KD. **(D)** Schematic representation of APD modeling in HAMs. **(E)** Population distribution of APD_90_ values for Control and Kmf 25% (=PLN KD) in HAMs (Left) and representative AP traces (right) showing the shortening effect on APD of simulated PLN KD.

In parallel, we screened 24 fly genes, orthologous to 17 of the 20 AF-associated genes. Genes were knocked down using a heart-specific driver (Hand-Gal4 (Sellin et al., 2006)) crossed to UAS-candidate gene-RNAi lines. Progeny of the crosses were aged to three-weeks old (middle aged) and heart function was characterized. Thirteen of the genes tested exhibited significantly altered systolic intervals and/or rhythm phenotypes in the fly cardiac model and 7 of these overlapped with the genes affecting APD in the ACMs (p-value < 0.001; **Fig. 3A**). In particular, cardiac-specific KD of KNCJ5/Irk3, GATA4-6/pnr, PITX2/Ptx1, and KCNA5/Sh resulted in prolonged SIs, consistent with the increased APD observed for ACMs. Cardiac KD of KCND3/Shal and PLN/SclA significantly shortened SIs, paralleling the reductions in APD observed in ACMs. Though no significant changes in arrhythmicity were observed in the ACMs, we did observe increased arrhythmicity (AI and MAD parameters) in flies in response to cardiac KD of three genes (Irk2, Pnr, and Sk; Wilcoxcon-ranked sum test, p-values < 0.001) in Drosophila.

Although there is evidence that APD prolongation is associated with AF (Nielsen et al., 2013; Olson et al., 2006), APD shortening is thought to be the most common mechanism underlying the onset and maintenance of AF (Teh et al., 2012; Wakili et al., 2011). We therefore focused on the gene KD that induced the strongest APD shortening phenotype. Remarkably, in both ACM and fly heart platforms, reduced PLN/SclA function consistently led to the strongest APD and SI shortening phenotype. In ACMs, PLN KD caused a significant shortening of median APD_75_ values, from 118.1 ms to 78.5 ms (∼ −40 ms) (KS-D=0.3305, p-value < 0.0001) and with calcium transient duration **(Fig. 3B and Suppl Fig. 3D**). Similarly, cardiac-specific KD of SclA (the PLN homolog in fly) significantly shortened contractions of the fly heart from 216 ms to 165 ms (**Fig. 3C)**.

To validate the phenotypic platform findings, we next employed a computational model of adult human atrial myocyte (HAM) (**Fig. 3D**) (Grandi et al., 2011). Here, we generated a population of 600 HAMs by randomly varying model parameters to replicate cell-to-cell variability (Morotti et al., 2017; Ni et al., 2018) and simulated PLN KD by increasing the affinity of the sarco/endoplasmic reticulum Ca2+-ATPase (SERCA) for cytosolic Ca2+ (i.e., the forward mode parameter Kmf was decreased to 25% of its control value, which mimics a strong PLN KD). Consistent with our phenotypic platform findings, our simulations showed significantly abbreviated APD90 (229 ms vs. 253 ms) (**Fig. 3E**) and Ca2+ transient duration (**Suppl Fig. 3E**) for PLN KD HAMs as compared to control groups paced at 2 Hz. Thus collectively, our multi-model system approach identifies PLN loss of function as conserved and most potent hit driving APD shortening among AF-associated genes.

### PLN loss of function-induced arrhythmia depends on β-adrenergic pathway stimulation and co-culture with fibroblasts

Next, to determine if loss of function of PLN alone is sufficient to induce arrhythmia-like phenotypes in ACMs, we measured the beat-to-beat interval variance (arrhythmia index, AI) in ACMs upon PLN KD, and found no difference as compared to siControl (**Suppl Fig. 4A-D)**. Additional factors such as conductance heterogeneity due to atrial fibrosis (Dzeshka et al., 2015; Xintarakou et al., 2020) as well as sympathetic stresses (Workman, 2010) have been tightly linked with the onset and maintenance of AF, and are often categorized as “AF-associated risk factors”. Thus, to mimic these environmental perturbagens, we co-cultured ACMs with fibroblasts for two days and/or applied the β-adrenoreceptor agonist, isoproterenol (1μM), acutely upon kinetic imaging. Remarkably, co-culturing ACMs with fibroblasts in a 3:1 ratio, nearly doubled the percentage of ACMs with AI values>20 (from 14.6% to 28.9%), however, in this context, PLN KD did not further increase the percentage of arrhythmic cells (**Suppl Fig. 4D-F)**. Conversely, treating ACMs with isoproterenol alone did not increase the percentage of arrhythmic cells, while exposing ACMs to Isoproterenol along with PLN KD, increased the percentage of arrhythmic cells from 13% to 22% (**Suppl Fig. 4G)**. Finally, co-culture with fibroblasts followed by acute isoproterenol treatment caused severe arrhythmia-like phenotypes (AI values > 20) in ∼ 38% of ACMs as compared to 20% in perturbagens only (**Fig.4A,B**). Interestingly, analysis of AP trains (lower panel of **Fig. 4B**) revealed missing beats and smaller refiring events that could be associated with delayed afterdepolarizations (DADs). Thus, collectively our results indicate that reduced PLN function predisposes ACMs to arrhythmia upon sensitization by fibroblasts and acute β-adrenergic stimulation.

**Figure 4.**
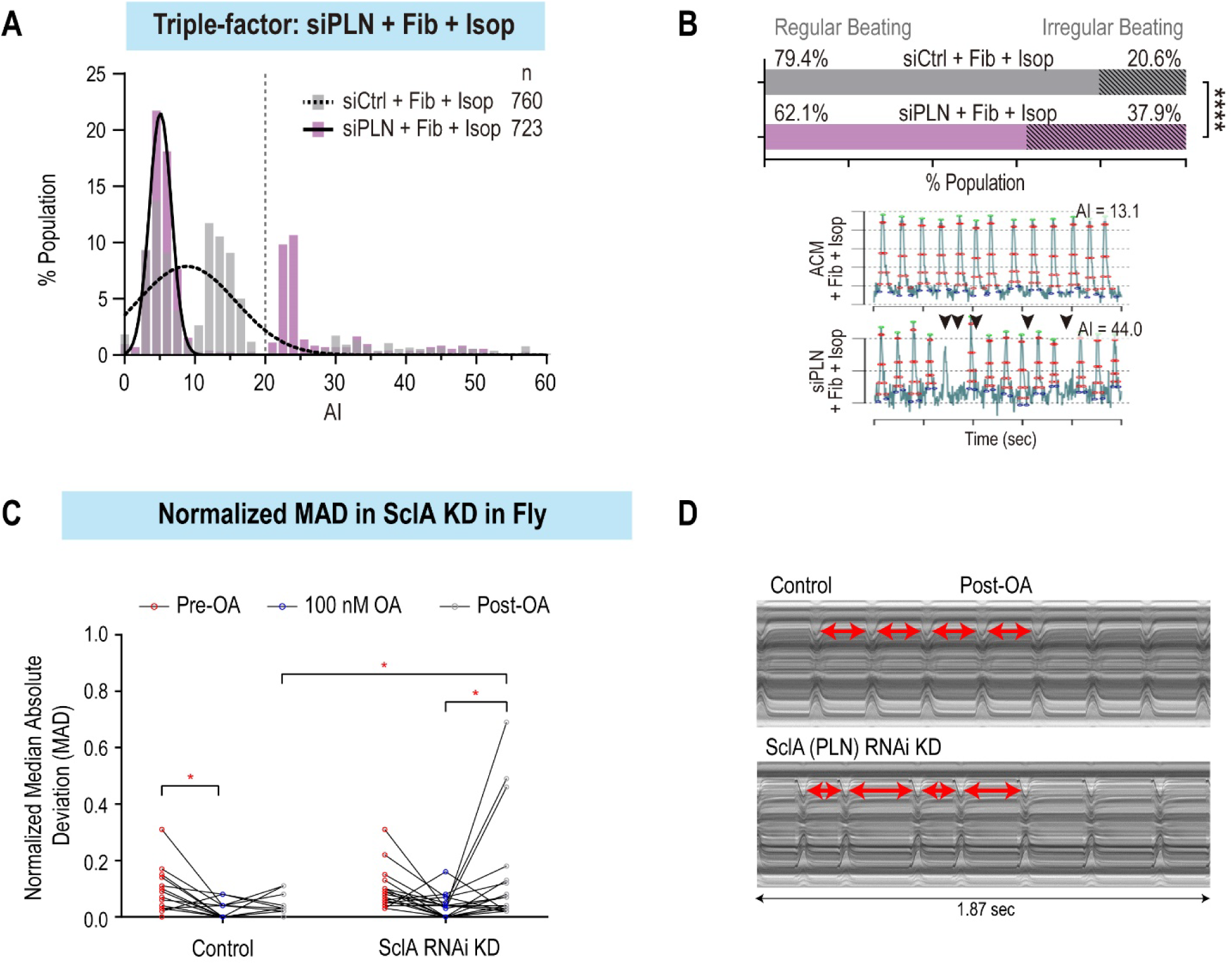
PLN KD induces arrhythmia phenotypes in combination with environmental pertubagens in both ACMs and flies. **(A)** Histogram of arrhythmia index (AI) values of ACMs co-cultured with fibroblasts and treated with isoproterenol (Isop) in siControl vs siPLN conditions. **(B)** Histogram showing the increased percentage of irregularly beating (AI>20) of ACMs co-cultured with fibroblasts and treated with Isop, in siPLN as compared to siControl. (Bottom) Representative peak trains of APs show irregular beat to beat interval (black arrowheads) in siPLN as compared to siControl condition. **(C)** Distribution of Median Absolute Deviation (MAD) values before, during and after 100nM octopamine treatment (OA). Post-OA, SclA KD hearts exhibit increased arrhythmia as compared to controls (p-value < 0.05, repeated measures 2-way ANOVA). **(D)** Representative M-modes showing irregular beat to beat intervals in SclA KD hearts post-OA as compared to control (Arrows show individual heart periods).

In fly hearts, despite significant changes in SI, neither AI nor MAD arrhythmia parameters were significantly altered by cardiac-specific PLN/SclA KD. To add an adrenergic stress, we exposed the fly hearts to octopamine (OA), the fruit fly version of norepinephrine/adrenaline (Sujkowski et al., 2017). Acute OA exposure significantly elevated heart rate by significantly shortening systolic intervals in both control and KD lines with a maximal effect at 100nM, which was the dose used for subsequent pharmacological pacing of the fly heart (**Suppl Fig. 5A,B)**. In controls, the mean SI returned to pre-exposure values at 10 min post-OA exposure (**Supp. Fig. 5C**), whereas the PLN/Sln KD hearts did not (**Supp. Fig. 5D**). In addition to SI, both contraction and relaxation intervals were significantly shortened in the presence of OA (**Supp. Fig. 5E,F**). We also observed increased post-octopamine pacing bouts of arrhythmia in the PLN/SclA KD hearts (mean nMAD = 0.1342; **Fig. 4C,D**) as compared to hearts from controls (mean nMAD= 0.0379; p-value: 0.019, repeated measures two-way ANOVA).

### Computational modeling validates PLN as a key regulator of rhythm in human adult atria

Next, to validate the phenotypic platform findings, we used models of both isolated HAMs and two-dimensional atrial tissue that allows to modulate cell-cell electrical coupling (**Fig. 5A** and (Colman et al., 2013; Ni et al., 2018)), that accounts for electrotonic effects of fibroblasts. In these assays, we applied a 2-Hz pacing-pause protocol to stimulate isolated HAMs or the left side of the atrial tissue construct and subsequently analyzed membrane voltage dynamics following the pause of pacing. We first tested the effect of increasing PLN KD on isolated HAMs, (Kmf 25% =high KD, Kmf 75% = low KD) along with simulated isoproterenol treatment. Remarkably, increasing PLN KD levels, led to the enhanced generation of triggered activity, which was further exacerbated by the simulated isoproterenol treatment (**Fig. 5B,C**). In this context, increasing PLN KD levels in combination with simulated isoproterenol treatment also led to an increase in occurrence of early afterdepolarizations (EADs) in isolated HAMs (**Fig 5C**, *bottom panel*). At the tissue level and similar to fibroblasts co-culture with ACMs, PLN KD induced more triggered activity (DADs and triggered APs) when combined with treatment with isoproterenol and reduced cell-cell electrical coupling (**Fig.5D-F, Suppl. Fig.6 A,B, and movies s1,2**). Collectively, our findings suggest that PLN loss of function predisposes cells to arrhythmia in a tissue environment with reduced electrical coupling and elevated β-adrenergic activity.

**Figure 5.**
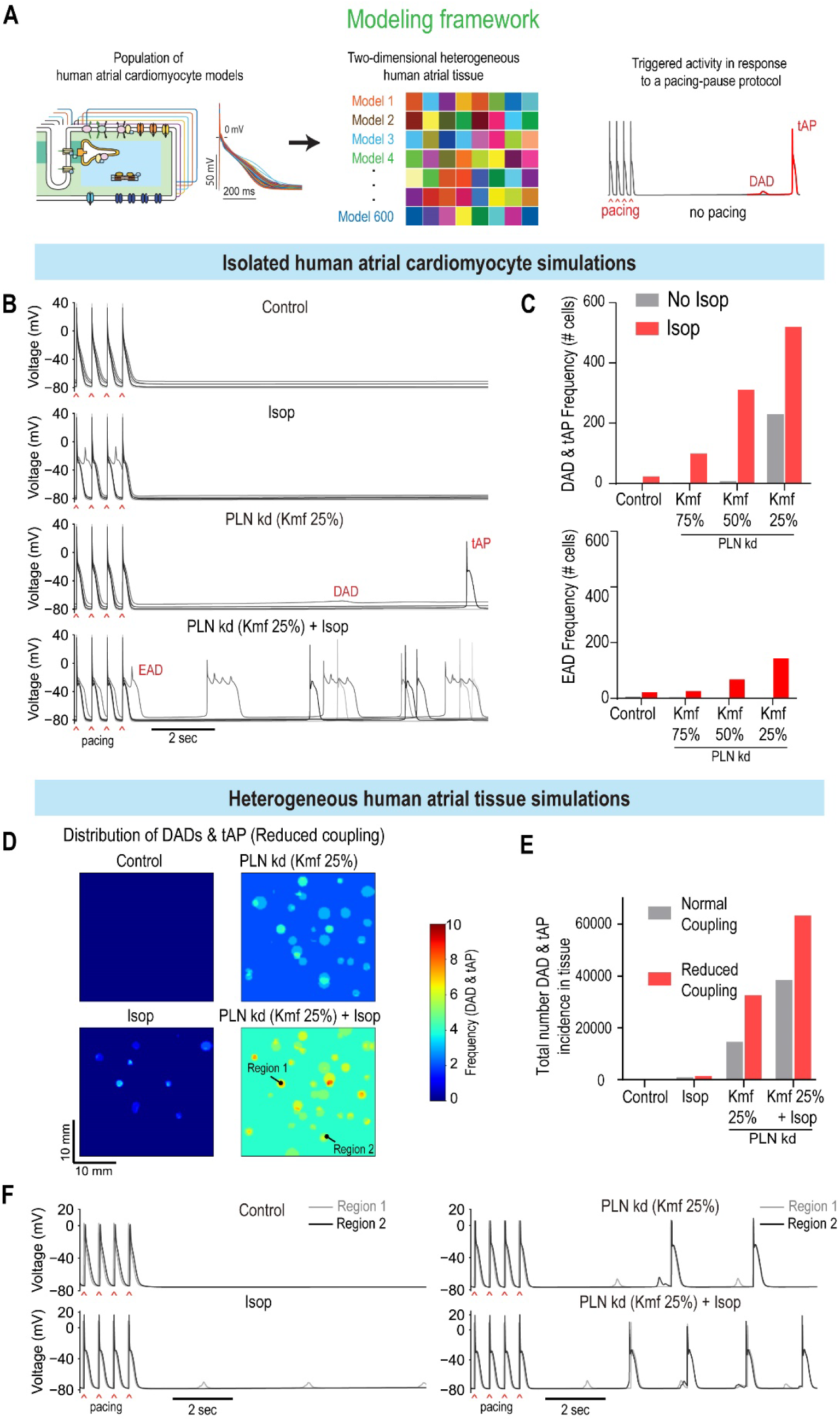
Combined PLN KD and Isop challenge promotes arrhythmic events in both isolated human atrial myocytes (HAMs) and two-dimensional atrial constructs. **(A)** Modeling framework for evaluating arrhythmic events in HAMs and two-dimensional (2D) human atrial tissue. For the 2D model of human atrial tissue, the physiological properties of each myocyte cluster were randomly assigned, thereby producing a heterogeneous tissue structure. A pacing (2 Hz)-pause protocol was applied to assess the incidence of triggered activities. **(B-C)** Effects of PLN kd (Kmf 25%) and Isop on the triggered activity in human atrial cardiomyocytes: **(B)** Time courses of APs for baseline (control), with Isop treatment, PLN KD (= Kmf 25%), and combined Isop treatment and PLN KD (PLN KD + Isop); **(C)** incidence of (top) DAD and tAP, and (bottom) EAD detected in the HAM populations for Isop and various degrees of PLN KD (Kmf varied from 25% to 75%) conditions. **(D-F)** Effects of PLN KD (Kmf 25%) and Isop on the generation of triggered activity in heterogeneous human atrial tissue: **(D)** spatial distribution of DADs and tAPs in the atrial tissue with reduced cell-to-cell coupling for PLN KD (Kmf 25%) and after Isop treatment; **(E)** total number of DADs and tAPs detected in the atrial tissue after each perturbation with normal or reduced cell-to-cell coupling; **(F)** Superimposed traces of APs from two regions (marked in panel *D*) of the atrial tissue with reduced cell-to-cell coupling for each perturbation.

### PLN functionally interacts with NCX and L-type calcium channels to control rhythm

To delineate aspects of PLN mode of action in the regulation of arrhythmia phenotypes, we next analyzed the mechanisms underlying the generation of DADs in HAMs. Strikingly, while at Kmf 25% (equivalent to 75% reduction in PLN) most cells generated DADs, whereas at Kmf 50%, only half of the cells generated DADs (see **Fig.5C**). To uncover the mechanism underlying DAD-generation in HAMs at Kmf50%, we separated DAD-generating HAMs from non-DAD-generating and applied logistic regression analysis (Morotti and Grandi, 2017) to determine the link between model parameters and DAD incidence (**Fig. 6A and Supplemental Table 2**). This analysis predicted that increased Ca^2+^ current I_CaB_, L-type Ca^2+^ current I_CaL_, or RyR release flux (i.e., by augmenting the parameters G_CaB_, G_CaL_ or V_RrRRel_) would promote the propensity for developing DADs. Similarly, our model also predicted that DAD occurrence correlated with reduced function of sodium-calcium exchanger NCX, RyR leakiness, sodium-potassium pump NaK, or ultra-rapid delayed rectifier K+ current I_Kur_ (**Fig. 6B**).

**Figure 6.**
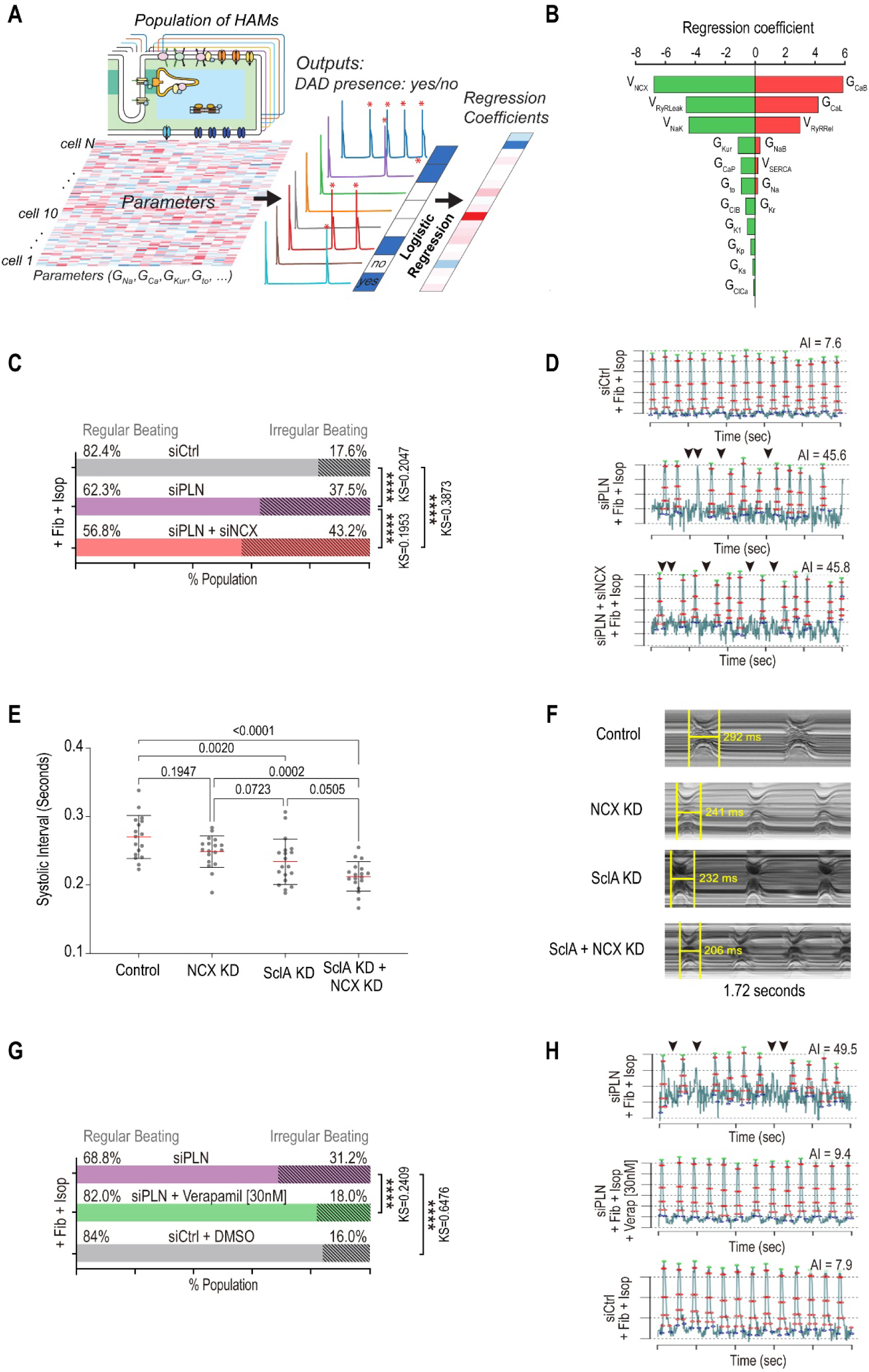
Multiple perturbations are required to generate arrhythmicity across platforms. **(A)** Schematic describing the logistic regression analysis approach to identify the mechanisms underlying the generation of DADs in HAMs. **(B)** Logistic regression analysis of DAD incidence in the context of moderate PLN knockdown (Kmf 50%) revealed influence of model parameters on the genesis of DADs in the population of HAMs in response to the pacing-pause protocol. Positive coefficients indicate that increasing the associated parameters promotes DAD production, and vice versa. **(C)** Histogram showing that the percentage of irregularly beating (AI > 20) ACMs co-cultured with fibroblasts and treated with isoproterenol, is increased when transfected with siPLN and NCX as compared to siPLN alone. **(D)** Representative AP peak trains for siControl, PLN siPLN +NCX conditions in ACMs co-cultured with fibroblasts and treated with isoproterenol. Arrowheads show examples of irregular beat to beat intervals in arrhythmically beating ACMs. **(E)** Mean Systolic Interval (SI) in response to cardiac KD of the plasma membrane Na^+^/Ca^2+^ exchanger NCX, SclA, and combined SclA + NCX KD. Co-KD caused a greater decrease in SI than did single KD alone (p < 0.05, Wilcoxon ranked sum test). **(F)** Representative m-modes showing effects of KD on SI. **(G)** Histogram showing that the percentage of irregularly beating (AI > 20) ACMs co-cultured with fibroblasts and treated with isoproterenol, is decreased when treated with verapamil (30nM) as compared to DMSO.

To validate these predictions, we selected two parameters that were most positively (ICaL conductance) or negatively (NCX maximal transport rate) correlated with DAD incidence. First, we tested in ACMs whether reduced expression of the sodium-calcium exchanger NCX, in the background of PLN KD would further increase the percentage arrhythmic cells. Consistent with the model prediction, combined KD of PLN and NCX in the presence of perturbagens (fibroblasts co-culture and isoproterenol infusion) significantly increased arrhythmia-like phenotypes as compared to single PLN KD, from 37.5% to 43.2% of cells with AI > 20 (**Fig. 6C**). Notably, the increased arrhythmic phenotypes of siPLN/NCX treated ACMs was specifically accompanied by very short APs as compared to siPLN alone (**Fig. 6D**). Next, to determine whether the PLN-NCX interaction is conserved at the whole organ level, we performed both single KD and co-KD of NCX//Calx and PLN/SclA using the Hand4.2-Gal4 heart-specific driver line flies. Remarkably, median SI was reduced in response to cardiac KD of SclA/Pln KD (232 ms) and by cardiac KD of NCX/Calx (253 ms, **Fig. 6E,F**). Co-KD of both genes further shortened the median SI to 211 ms (p = 0.05, Wilcoxon ranked sum test).

Finally, the regression analysis also revealed that DAD-generating HAMs had increased L-type calcium channel (G_CaL_) currents. Thus, to test whether inhibition of L-type calcium channels activity might reduce PLN-induced arrhythmia, we treated ACMs with a calcium channel blocker, verapamil, and quantified the percentage of arrhythmic cells in response to PLN KD+Fib+isoproterenol treatment. Remarkably, ACMs treated with verapamil were 1.7 fold less arrhythmic than DMSO control (from 31.2% to 18%, **Fig. 6G,H**). Collectively, our results indicate that PLN KD-dependent arrhythmia phenotypes are at least in part driven by NCX and L-type calcium channel activity. In addition, our results also highlight that the combinatorial use of computational modeling and phenotypic platforms represents a novel approach to identify novel gene interactions involved in the regulation of atrial rhythm and to predict potential therapeutic targets to treat AF.

## DISCUSSION

### 1) Multi-system platform to model AF

As for other common cardiovascular diseases (i.e hypertension, myocardial infarction), AF is caused by complex and mostly unknown combinations of genetic (Roselli et al., 2020) and environmental (i.e fibrosis, age)(Schuttler et al., 2020) insults, that render experimental modeling a challenging process. In this context, and due to the growing list of AF-associated genes (Christophersen et al., 2017; Nielsen et al., 2018a; Roselli et al., 2018), we assessed that an ideal modeling strategy should include the ability to rapidly test the role of large cohort of genes on cardiac electrophysiological and rhythm parameters, along with their interaction with other genes and/or environmental perturbagens (*i.e* age, fibrosis or hormonal stress), preferably in a human adult atrial context. To address some of these modeling requirements, we have assembled (Schuttler et al., 2020) a new phenotypic platform that enables rapid evaluation of gene function on the regulation of APD and rhythm in model systems providing *1)* a human and atrial context (ACM model), and *2)* an intact, functionally mature and aging heart (fly model). Because both ACM and fly systems have genome-wide screening capacity (Neely et al., 2010; Nielsen et al., 2022), they unlock the exploratory power of functional genomics that is needed for the unbiased identification of novel genes and pathways controlling cardiac rhythm beyond those identified in GWAS studies. A remarkable feature of the ACM platform is the ability to assess APD and rhythm parameters with single cell resolution, which enables *1)* the development of co-culture conditions mimicking aspects of known AF-associated risk factors such as fibrosis and *2)* the quantification of the heterogeneity in cellular responses to environmental perturbations. Similarly, in flies, rapid aging (5 weeks fly = old fly)(Blice-Baum et al., 2019; Ocorr et al., 2007a) and the ability to expose these animals to environmental stressors such as octopamine or different diets (high fat or high sugar) (Birse et al., 2010; Johnson et al., 1997), enables the study of genes’ function and their interaction with the environment in a physiologically integrated heart system. Because ACMs represent a relatively immature state of human adult atrial myocytes (Uosaki et al., 2015) and the fly heart architecture differs significantly from that of mammals, findings using these platforms need to be further validated in models with human adult atrial relevance. Thus, to address these limitations, we have incorporated computational models of human adult atrial myocytes as a third model and a tool for both validation (**Fig. 3 and 5**) and hypothesis generation (**Fig. 6**). In sum, we propose that the integrated use of model systems combining functional screening capacity and human atrial and whole organ physiological relevance, represent a novel approach enabling the identification and characterization of new genes affecting AF-associated rhythm biology with unprecedented throughput.

### 2) Cellular arrhythmia is a compound phenotype caused by multiple perturbagens

Cellular arrhythmias associated with AF are known to arise from “vulnerable substrates” caused by either APD prolongation or shortening, and thereby promoting early and/or delayed after-depolarizations or circuit re-entry, respectively. In this context, conductance heterogeneity due to atrial fibrosis, as well as sympathetic stresses, have been tightly linked with the onset of AF, and are often categorized as “AF-associated risk factors” (Fatkin et al., 2017). Here, we have replicated the effects of some of these risk factors (isoproterenol and/or fibroblasts in ACMs and octopamine in the fly heart) and observed that multiple stressors were required to produce robust arrhythmia phenotypes upon AF-associated gene KD (PLN). These results also emphasize the requirement for model systems that permit the incorporation of such risk factors for efficient arrhythmia modeling. In this context, a remarkable contribution of the fly model is the observation that cardiac arrhythmias can occur in response to ion channel KO and KD (Ocorr et al., 2007b; Ocorr et al., 2017). Interestingly, these arrhythmias only became robust with increasing fly age, and in the case of the KCNQ channel, this was linked to an age-related reduction in channel expression that could be reversed by channel overexpression in old flies (Nishimura et al., 2011). In sum, these new models facilitate the molecular delineation of how environmental insults synergize with genetic predispositions to induce arrhythmia-associated phenotypes and thus represent a promising new research direction to uncover novel therapeutic avenues to treat AF.

### 3) Mechanism of PLN-driven arrhythmia in atrial myocytes

In this study, we identified PLN loss of function as top hit, among the 20 AF-associated genes tested, causing both APD shortening and arrhythmia phenotypes (i.e beat to beat interval irregularities, DADs) when combined with environmental perturbagens. Consistent with a potential role for PLN as a AF-contributing gene, three large independent GWAS studies (Christophersen et al., 2017; Nielsen et al., 2018a; Roselli et al., 2018) have identified SNPs in the vicinity of PLN gene locus in patients with AF(Christophersen et al., 2017; Nielsen et al., 2018a; Roselli et al., 2018), although the functional significance of such variants is unknown. Mechanistically, our simulations in HAMs suggest that PLN loss of function selectively increases atrial SERCA pump activity and the sarcoplasmic reticulum Ca^2+^ load while shortening Ca^2+^ transient duration. The associated Ca^2+^overload and spontaneous, in turn, contribute to cause electrical instabilities. Consistent with our model, increased SERCA function is observed during paroxysmal AF (Denham et al., 2018; Voigt et al., 2014) and overexpression of SERCA in mouse atria promotes cellular correlates of AF (Nassal et al., 2015). Interestingly, similar results have also been reported with the ablation of PLN functional homolog, sarcolipin, where marked structural and electrical atrial remodeling was reported in mice (Babu et al., 2007). Furthermore, our computational modeling approach also indicated that PLN KD-induced arrythmia phenotypes are exacerbated by increased L-type Ca^2+^ current and decreased Na^+^/Ca^2+^ exchanger (NCX) activity. These simulations suggest that L-type Ca^2+^ channels (LTCCs) and NCX modify PLN activity and contribute to regulate rhythm in atrial myocytes and thus might represent a class of second hits contributing to promote PLN KD-induced arrythmias. Consistent with these predictions, co-KD of PLN and NCX in ACMs increased the percentage of cells displaying arrhythmic phenotypes while inhibition of L-Type calcium channel activity drastically reduced these phenotypes. In the fly heart, interactions between these same two genes were also pro-arrhythmogenic in that co-KD significantly shortened the SI relative to individual gene KD. Collectively, these observations suggest a central role for the PLN-SERCA-LTCCs-NCX axis in the regulation of AF-associated rhythm phenotypes, which is consistent with the mechanisms known to contribute to AF pathophysiology (reviewed in (Denham et al., 2018)), and thereby highlight the physiological relevance of this new approach to model AF.

### What’s next – Next steps

This platform provides an in-depth resolution of cardiac electrophysiology metrics with various applications: (1) large-scale functional genomic screens to identify novel gene regulatory networks governing cardiac rhythm; (2) establishment of new arrhythmia models to phenotypically characterize rhythm-associated cardiac diseases; (3) small molecule screens for anti-arrhythmic drug discovery.

### Limitation of the system

Limitations of the presented platform includes the use of hPSC-derived ACMs to study AF, a known age-dependent disease. Although cardiomyocyte differentiation techniques are advancing, single-cell RNA-seq results from various studies implied that hPSC-derived CMs are in similar transcriptional stages as pre-neonatal or fetal cardiomyocytes (DeLaughter et al 2016, Friedman et al 2018). Physiology differences, such as cellular morphometry, functional maturity, transcription profile, and disease manifestation, of these relatively immature hPSC-derived ACMs may undermine the fidelity to study a highly age-dependent disease. To mimic the conductance heterogeneity due to cardiac fibrosis, fibroblasts were co-cultured with the ACMs in a mono-layer format. This simplified reproduction of in vitro cardiac fibrosis model may not be sufficient to fully recapitulate complex arrhythmia mechanisms that occur at the organ level. Such limitation may be overcome with the advancement of current culturing technics, including micropatterning cell culture substrates or cardiac organoids (Salick et al., 2014; Zhao et al., 2021). The lack of stimulus apparatus of the platform restricts certain pacing-induced arrhythmia studies. However, by incorporating compounds with beat rate regulating properties, such as β-adrenergic modulators in this study may provide an alternative approach of bradycardia/tachycardia-induced arrhythmias. Lastly, the siRNA knockdown approach applied in this study is limited in evaluating the loss of function of the AF-associated gene candidates. Previous studies indicate that the gain of function mutation (i.e., KCNA5) and the abnormally elevated expression level (i.e., PITX2), can contribute to the initiation and maintenance of AF (Christophersen et al 2012, Hernadez et al 2016). This can easily be tested in the Drosophila model, where tissue-specific gene overexpression is achieved using the same Gal4 driver system used for gene KD. We are currently developing an ACM screening assay that incorporates CRISPR-activation (CRISPRa) technique for the gain of function mutations screens (cite CRISPRa).

## MATERIAL AND METHODS

### Generation of ACMs

Id1 overexpressing hiPSCs^1^ were dissociated with 0.5 mM EDTA (ThermoFisher Scientific) in PBS without CaCl^2^ and MgCl^2^ (Corning) for 7 min at room temperature. hiPSC were resuspended in mTeSR-1 media (StemCell Technologies) supplemented with 2 µM Thiazovivin (StemCell Technologies) and plated in a Matrigel-coated 12-well plate at a density of 3 x 10^5^ cells per well. After 24 hours after passage, cells were fed daily with mTeSR-1 media (without Thiazovivin) for 3-5 days until they reached ≥ 90% confluence to begin differentiation. hiPSC-ACMs were differentiated as previously described^2^. At day 0, cells were treated with 6 µM CHIR99021 (Selleck Chemicals) in S12 media^3^ for 48 hours. At day 2, cells were treated with 2 µM Wnt-C59 (Selleck Chemicals), an inhibitor of WNT pathway, in S12 media. 48 hours later (at day 4), S12 media is fully changed. At day 5, cells were dissociated with TrypLE Express (Gibco) for 2 min and blocked with RPMI (Gibco) supplemented with 10% FBS (Omega Scientific). Cells were resuspended in S12 media supplemented with 4 mg/L Recombinant Human Insulin (Gibco) (S12+ media), 300 nM retinoic acid (R2625-50MG) and 2 µM Thiazovivin and plated onto a Matrigel-coated 12-well plate at a density of 9 x 10^5^ cells per well. S12+ media was changed at day 8 and replaced at day 10 with RPMI (Gibco) media supplemented with 213 µg/µL L-ascorbic acid (Sigma), 500 mg/L BSA-FV (Gibco), 0.5 mM L-carnitine (Sigma) and 8 g/L AlbuMAX Lipid-Rich BSA (Gibco)(CM media). Typically, hiPSC-ACMs start to beat around day 9-10. At day 15, cells were purified with lactate media (RPMI without glucose, 213 µg/µL L-ascorbic acid, 500 mg/L BSA-FV and 8 mM Sodium-DL-Lactate (Sigma)), for 4 days. At day 19, media was replaced with CM media.

### Voltage Assay in ACMs

Voltage assay is performed using the labeling protocol described in McKeithan et al., 20174. Briefly, hiPSC-ACMs at day 25 of differentiation were dissociated with TrypLE Select 10X for up to 10 min and the action of TrypLE was neutralized with RPMI supplemented with 10% FBS. Cells were resuspended in RPMI with 2% KOSR (Gibco) and 2% B27 50X with vitamin A (Life Technologies) supplemented with 2 µM Thiazovivin and plated at a density of 6,000 cells per well in a Matrigel-coated 384-well plate. hiPSC-ACMs were then transfected with siRNAs directed against AFib-associated genes (ON-TARGETplus Human, siGATA4: J-008244-05-0002, siGATA5: J-010324-06-0005, siGATA6: J-008351-06-0005, siGJA1: J-011042-05-0002, siGJA5: J-017368-05-0002, siHAND2: J-008698-06-0005, siHCN4: J-006203-05-0002, siKCNA5: J-006215-06-0005, siKCND3: L-006226-00-0005, siKCNJ5: J-006250-06-0002, siKCNN3: J-006270-06-0002, siNKX2-5: J-019795-07-0002, siNKX2-6: J-025793-17-0002, siNPPA: J-012729-05-0002, siPITX2: J-017315-05-0005, siPLN: J-011754-05-0005, siSH3PXD2A: J-006657-07-0002, siSYNE2: J-019259-09-0002, siTBX5: J-013410-05-0002, siZFHX3: J-015412-05-0002) using lipofectamine RNAi Max (ThermoFisher). Each siRNA was tested individually in quadruplicates. Three days post-transfection, cells were first washed with pre-warmed Tyrode’s solution (Sigma) by removing 50 µL of media and adding 50 µL of Tyrode’s solution 5 times using a 16-channel pipette. After the fifth wash, 50 µL of 2x dye solution consisting in voltage-sensitive dye Vf2.1 Cl (Fluovolt, 1:2000, ThermoFisher) diluted in Tyrode’s solution supplemented with 1 µL of 10% Pluronic F127 (diluted in water, ThermoFisher) and 20 µg/mL Hoescht 33258 (diluted in water, ThermoFisher) was added to each well. The plate was placed back in the 37°C 5% CO_2_ incubator for 45 min. After incubation time, cells were washed 4 times with fresh pre-warmed Tyrode’s solution using the same method described above. hiPSC-ACMs were then automatically imaged with ImageXpress Micro XLS microscope at an acquisition frequency of 100 Hz for a duration of 5 sec with an excitation wavelength of 485/20 nm and emission filter 525/30 nm. A single image of Hoescht was acquired before the time series. Fluorescence over time quantification and trace analysis were automatically quantified using custom software packages developed by Molecular Devices and Colas lab. At least, two independent experiments were performed.

### Arrhythmia Assay and Drug Treatment in ACMs

hiPSC-ACMs were dissociated, plated in 384-well plate and transfected with siRNA-associated AFib as described previously (ON-TARGETplus Human, siNCX: J-007620-05-0002). 24 hours post-transfection, 2,000 primary human fibroblasts were added per well to the hiPSC-ACMs. 48 hours later (the day of the imaging), cells were dyed with the voltage sensitive dye Vf2.1 Cl as described above, then treated with 50 µL of 2x solution of isoproterenol (1 µM final) diluted in Tyrode alone and in combination with 2x solution of Verapamil (30 nM final), diluted in Tyrode at the 5^th^ wash. After 20 minutes of compound incubation time, cells were imaged, and single-cell traces analyzed as described previously.

### Whole Cell Patch Clamp Electrophysiology

Cardiac ion currents were recorded from single cardiomyocytes using the whole-cell patch-clamp method. Briefly, coverslips with ACMs or VCMs were transferred into electrophysiological perfused recording chamber (RC-25-F, Warner Instruments, Hamden, CT) mounted on the stage of an inverted Olympus microscope. Patch pipettes were pulled from thin-wall borosilicate glass capillaries (CORNING 7740, 1.65mm) with a P-2000 laser pipette puller (Sutter Instruments, California, USA) and had electrode tip resistances between 1.5 and 5.5 MΩ with access resistance of <8MΩ for whole-cell patch recordings. Series resistance and cell capacitance were compensated to between 30 and 60% in some voltage-clamp recordings. Patch electrodes were filled with intracellular solution containing: 120mM CsCl, 20mM tetraethylammonium chloride (TEA-Cl), 10mM Hepes, 2.25mM EGTA, 1mM CaCl2, 2mM MgCl2, pH 7.4. All recordings were performed at room temperature in Tyrode’s solution. Current response traces were acquired using the Axon 200B amplifier. Currents were digitally sampled at 10 kHz using Digidata 1322A digitizer hardware and pClamp 10.2 software (Molecular Devices, California, USA). For both ACMs and VCMs, n=5.

### Drosophila Strains

We used the Hand 4.2-Gal4 fly line as our heart-specific driver line (Brand and Perrimon, 1993). Virgin Hand-Gal4, females were crossed to male flies from UAS-RNAi lines for each AF gene candidate. UAS-RNAi lines and their respective control lines were acquired from the Bloomington Drosophila Stock Center (BDSC, Indiana, United Sates of America) and Vienna Drosophila Resource Center (VDRC, Vienna, Austria). For each gene candidate, at least 2 different RNAi lines were used (see **Supplemental Table 2**; GD and KK were the genetic background lines for stocks from VDRC lines and ATTP2 and ATTP40 were the genetic background lines for stocks from BDSC).

The PLN (fly ortholog: Sarcolamban A / SclA) sensitized fly line was made by recombining the USA-SclA RNAi with the Hand 4.2-Gal4 heart-specific driver line (Kaplan and Trout, 1969; Jan et. al, 1977; Gardnell et al 2006). Virgin females from the Hand-Gal4 or the SclA-sensitized, Hand-Gal4 driver lines were crossed to males of the desired UAS-RNAi lines. Adult female flies for all crosses were collected upon eclosion and raised at 25°C on a 12-hour light-dark cycle. Flies were fed a standard yeast-cornmeal diet, with food replaced every other day.

### Drosophila Heart Function Characterization

Cardiac phenotypes of middle-aged (3-weeks old) female flies from each cross were characterized using denervated, semi-intact preparations as previously described in (Ocorr et al., 2009; Vogler and Ocorr, 2009). Briefly, hearts from 20-25 flies were examined for each genotype and age. Adult female flies were exposed to FlyNap, a triethylamine-based anesthetic, for at least one minute until no movement was detected. Hearts were exposed by dissection in room temperature, air bubbled, artificial hemolymph (AHL, Ocorr et al, 2007). High-speed video recordings were filmed with a Hamamatsu EM-CCD camera and using HC Image capture software (Hamamatsu Corp). Heart movements were analyzed using the Semi-automated Optical Heartbeat Analysis (SOHA) software (http://sohasoftware.com/). Movies were recorded at speeds of 140+ fps with pixel resolution ∼ 1 micron/pixel allowing very precise temporal and spatial measurements, including heart period (HP) and rate (1/HP), diastolic and systolic intervals, and fractional shortening/contractility. To quantitate arrhythmia, we first calculate the median absolute deviation (MAD). The median value of the absolute deviations of each heart period (Xi) from the median heart period 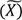 is calculated and then multiplied by a constant (k = 1.4826 assuming data is normally distributed).

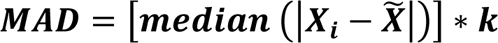

To normalize the MAD index (nMAD), the MAD value was divided by the median heart period. Qualitative records of heart wall movements (M-modes / kymographs) were produced by electronically excising a 1 pixel horizontal “slice” from each movie frame and aligning them horizontally providing an edge trace displaying heart wall movements in the X-axis over time along the Y-axis (Ocorr et al., 2009; Ocorr et al., 2007b).

### Octopamine-challenge Heart Assay

Octopamine (OA) pacing experiments were performed in situ on the semi-intact fly preparation. OA (Sigma-Aldrich #O0250) stock solution (10 mM) was freshly prepared by dissolving in water and was further diluted in AHL. A dose-response curve was generated using doses ranging from 0.1 nM OA to 500 nM OA (Supplemental Fig. 5A). The increase in heart rate was maximal at 100 nM OA, which was the dose used for all subsequent pacing experiments (Supplemental Fig. 5B). Following dissection, hearts were first allowed to equilibrate in fresh AHL for 15 min and 30 sec movies of heart function were recorded. Heart function was recorded three times per fly: 1) pre-drug exposure, 2) after a 15-minute exposure to 100nM OA, and 3) after a 15-minute post drug exposure recovery period. A second set of hearts exposed only to vehicle (AHL) were filmed at the same three 15-minute intervals to serve as time controls.

### Simultaneous optical and electrophysiological recordings

Simultaneous optical and intracellular electrical recordings were performed as previously described in (Ocorr et al., 2017). Briefly, we used a semi-intact preparation that was incubated in artificial hemolymph. Optical recordings were done as described above; electrical potentials were recorded using sharp glass electrodes (20±50MΩ) filled with 3M KCl and standard intracellular electrophysiological techniques. Data were acquired using an Axon-700B Multiclamp amplifier, signals were digitized using the DIGIDATA 1322A and data were captured and analyzed using PClamp 9.0 and Clampfit 10.0 software respectively (all from Molecular Devices). Data was quantified from representative 30s recordings where the resting membrane potential had remained stable for at least 30s. To coordinate the optical and electrical recordings a TTL pulse was sent by the image capture software to the Digitizer. The pulse duration lasted for the entire period of optical recording and was recorded in a separate channel by the PClamp software allowing us to delineate the beginning and the end of the optical recording and directly align it with the electrical record.

### Statistical Analysis

ACMs - Population distribution of control and siRNA-treated hiPSC-ACMs was generated with GraphPad Prism software (2019) using nonlinear regression. Unpaired nonparametric Kolmogorov-Smirnov test was used to compare each treated conditions to control using APD_75_ of every measured cells. To determine any statistical significance between experimental and control groups, we calculated two-sided p-values with Student’s t-test using GraphPad Prism software.

Flies **-** Data that exhibited a normal distribution was evaluated for significance using a 1-way ANOVA (for simple comparisons) or a 2-way ANOVA (for multiple manipulations) followed by multiple comparisons post-hoc tests as indicated in figure legends. Data sets that did not show a normal distribution (typically heart period, systolic interval and diastolic interval) were analyzed using a nonparametric Wilcoxon Rank Sum test or Kruskal-Wallis test followed by Dunn multiple comparisons post-hoc tests. For acute octopamine stress experiments, we used repeated measures, 2-way ANOVA followed by Sidak’s multiple comparisons test. Statistical analysis and data visualization was completed with GraphPad Prism (v8.0.0; www.graphpad.com), R (v3.6; https://www.r-project.org), and Rstudio (v1.3.959; https://rstudio.com).

### Computational Modeling Design

We employed our well-established computational model (Grandi et al., 2011) of human atrial myocytes to simulate human action potential (AP) and Ca2+. PLN regulates the function of sarco/endoplasmic reticulum Ca2+-ATPase (SERCA) function by decreasing the apparent affinity of SERCA for Ca2+ ions (Periasamy et al., 2008; Simmerman and Jones, 1998). Accordingly, the effects of PLN knockdown on SERCA were simulated by various degrees of reduction in the SERCA affinity parameter (Kmf) for cytosolic Ca2+: Kmf was scaled by 75%, 50%, or 25% to cover a wide parameter space of change. These changes were made based on a previous study showing that applying PLN antibody shifted the affinity from 0.8 μM to 0.2 μM (Cantilina et al., 1993).

### Modeling Arrhythmias in Human Atrial Cells

To describe the intrinsic cell-to-cell variabilities in atrial electrophysiology and uncover the uncertainty of the modeling results, we applied a population-based approach (Ni et al., 2018; Sobie, 2009) built populations of 600 human atrial model variants by randomly perturbing key model parameters (e.g., the maximum ion channel conductances, rates for membrane transporters, and Ca2+ handling fluxes; detailed in **Table 1**) by a lognormal distribution (σ = 0.2).

### Logistic Regression Analysis of Delayed afterdepolarizations

We performed logistic regression analysis (Morotti et al., 2017) to understand the influence of each model parameter on the arrhythmic outcome in human atrial myocytes. For each cell of the population of models, a binary code (yes/no) was applied to describe the presence/absence of delayed afterdepolarizations (DADs). Logistic regression coefficients were obtained using MATLAB (R2019b) scripts as detailed previously in Morotti and Grandi, 2017.

### Modeling Arrhythmias in Human Atria Tissue

We created two-dimensional (2D) models to understand the dynamic behaviors of atrial AP and Ca2+ in tissue using monodomain equations to describe the tissue electrical coupling as did in our previous studies (Ni et al., 2017). The 2D model comprises 120 × 125 grids with a spatial interval of 0.3 mm. To account for the intrinsic variabilities in tissue, we mapped our population of models to the tissue based on a heterogeneous pattern dividing the tissue into 600 blocks consisting of 5×5 grids. To assess how tissue coupling affects the arrhythmic events, we also simulated a reduced coupling (scale to 25% of tissue conductivity) condition.

### Pacing-and-pause protocol in single cell and tissue stimulation

A constant pacing-and-pause protocol was applied to evaluate the physiological effects of PLN knockdown. Specifically, single cells were paced at 2 Hz for 290 s prior to a 10-s period of pause without stimulation. In tissue simulations, stimuli were applied at the left side of the 2D tissue at 2 Hz for 10 s, which was followed by a 10-s period without stimulation. AP and Ca2+ traces from the last four stimuli and the non-paced period were recorded for data analysis. Logistic regression analysis was applied to uncover the influence of model parameters on the incidence of arrhythmogenic events.

## Supporting information

All supplemental Figures

## ACKNOWLEDGEMENTS

This work was supported by grants DISC2-10110 (California Institute for Regenerative Medicine), R01 HL153645, R01 HL148827, R01 HL149992, R01 AG071464 (National Institutes of Health), and SBP institutional support to AC. NIH R01 HL13224 supported KO and F32 HL56607 J.K.

---

**Supplemental Figure 1.**
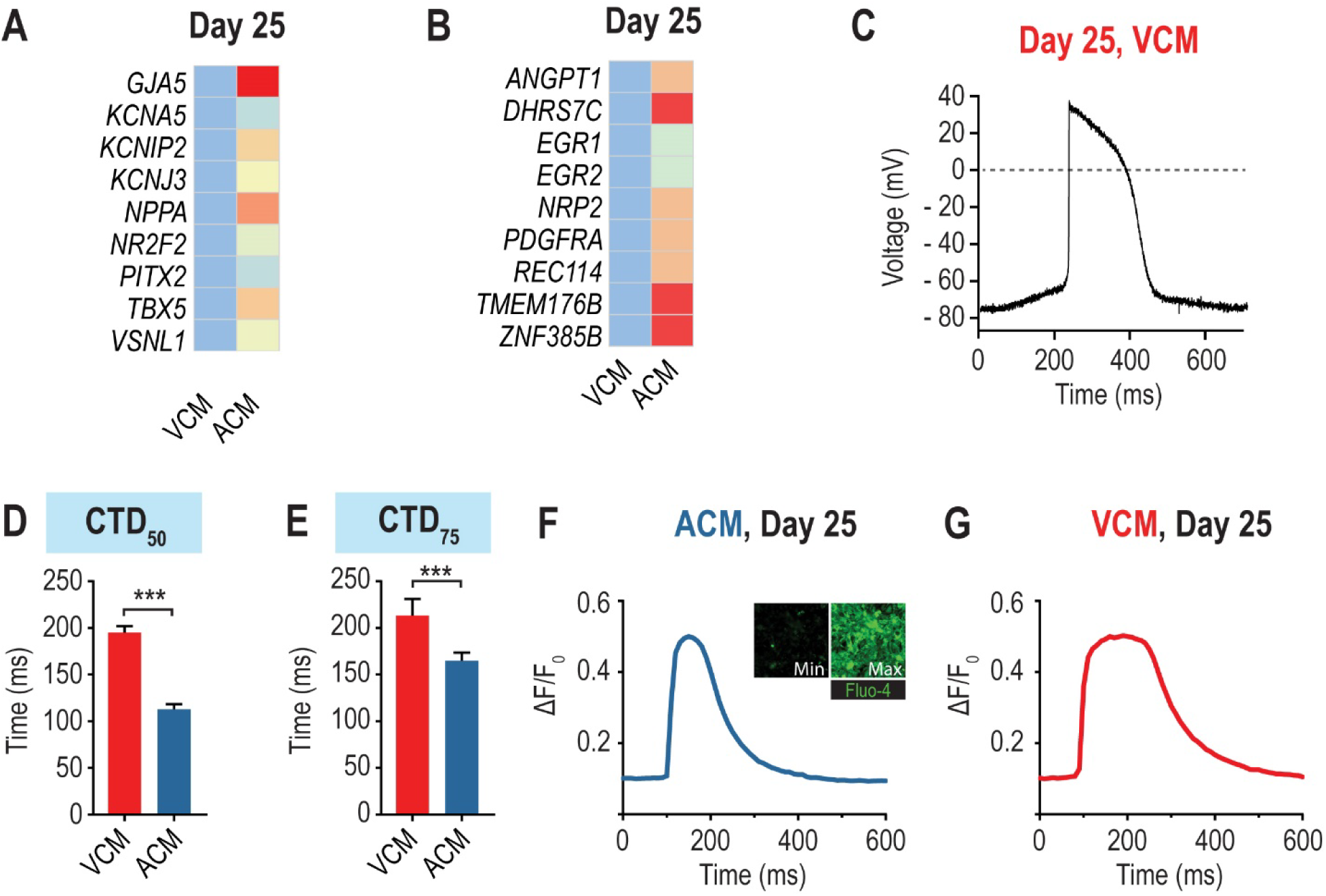
Gene expression and functional characterization of ACMs. **(A & B)** Heatmap of atrial genes enriched in day 25 ACMs as compared to VCMs. **(C)** Patch-clamp experiments show that VCMs generate action potential with ventricular characteristics, including a long phase 2. **(D,E)** Quantification of average CTD50 and CTD75 demonstrate that ACMs display shorter calcium transients than VCMs. P-value *** < 0.001. **(F,G)** Representative calcium transient traces for ACMs and VCMs respectively.

**Supplemental Figure 2.**
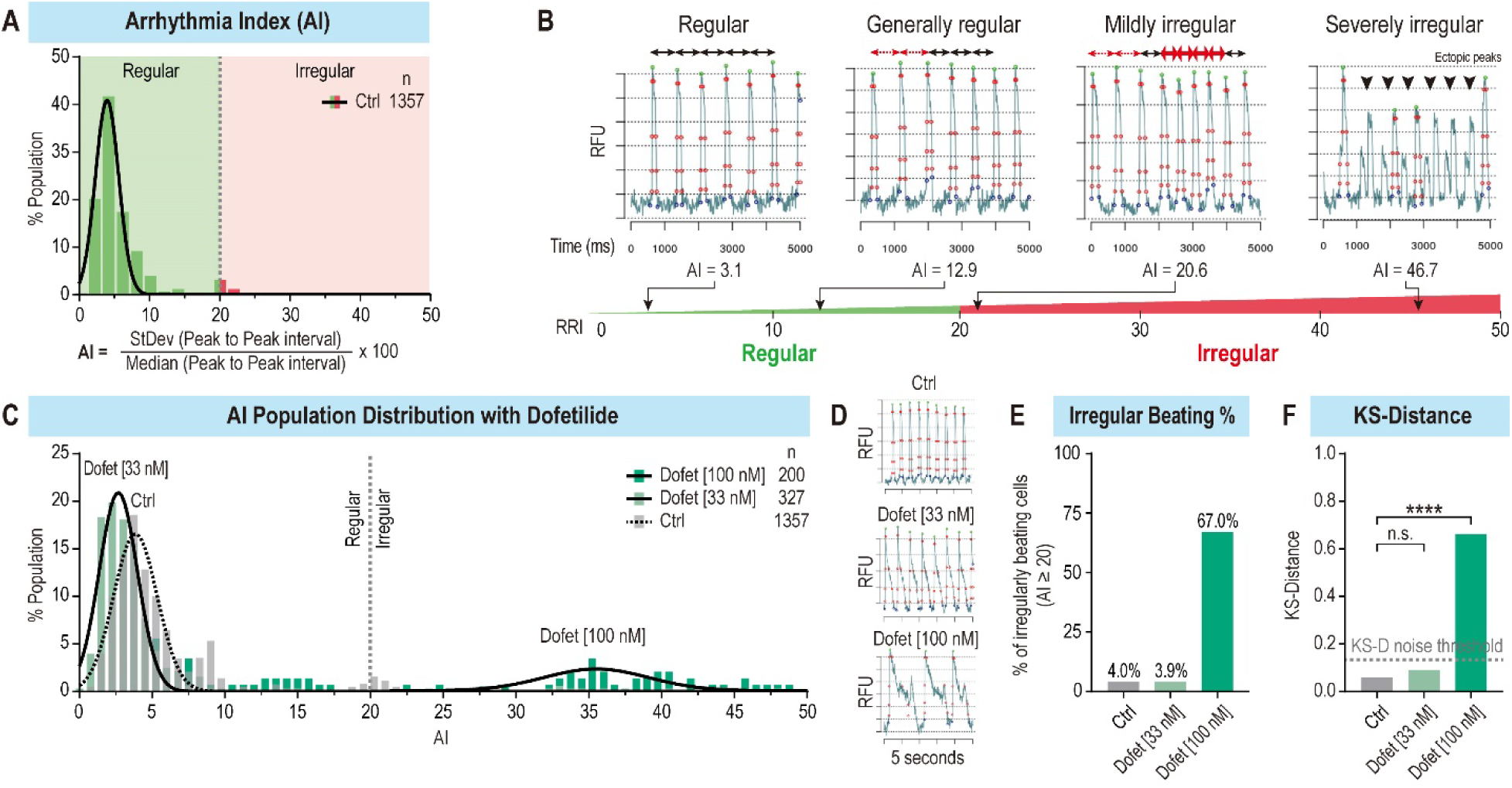
Quantification of rhythm parameters in ACMs. **(A)** Example of histogram with Arrhythmia Index (AI) values showing cells with AI<20: not arrhythmic or AI > 20 arrhythmic. **(B)** Examples of peak trains from ACMs with regular (left), mildly irregular (middle), and severely irregular (right) with their respective AI score. **(C)** Histogram representing the distribution of AI values for ACMs treated with increasing doses of Dofetilide. **(D)** Representative peak trains of control ACMs (top), ACMs treated with 33 nM Dofetilide (middle), and ACMs treated with 100 nM Dofetilide (bottom). **(E)** Histogram representing the proportion of arrhythmic cells (AI > 20) in control ACMs and ACMs treated with 33 nM and 100 nM Dofetilide. **(F)** Quantification of KS-D between control or Dofetilide-treated ACMs conditions. P-value *** < 0.0001.

**Supplemental Figure 3.**
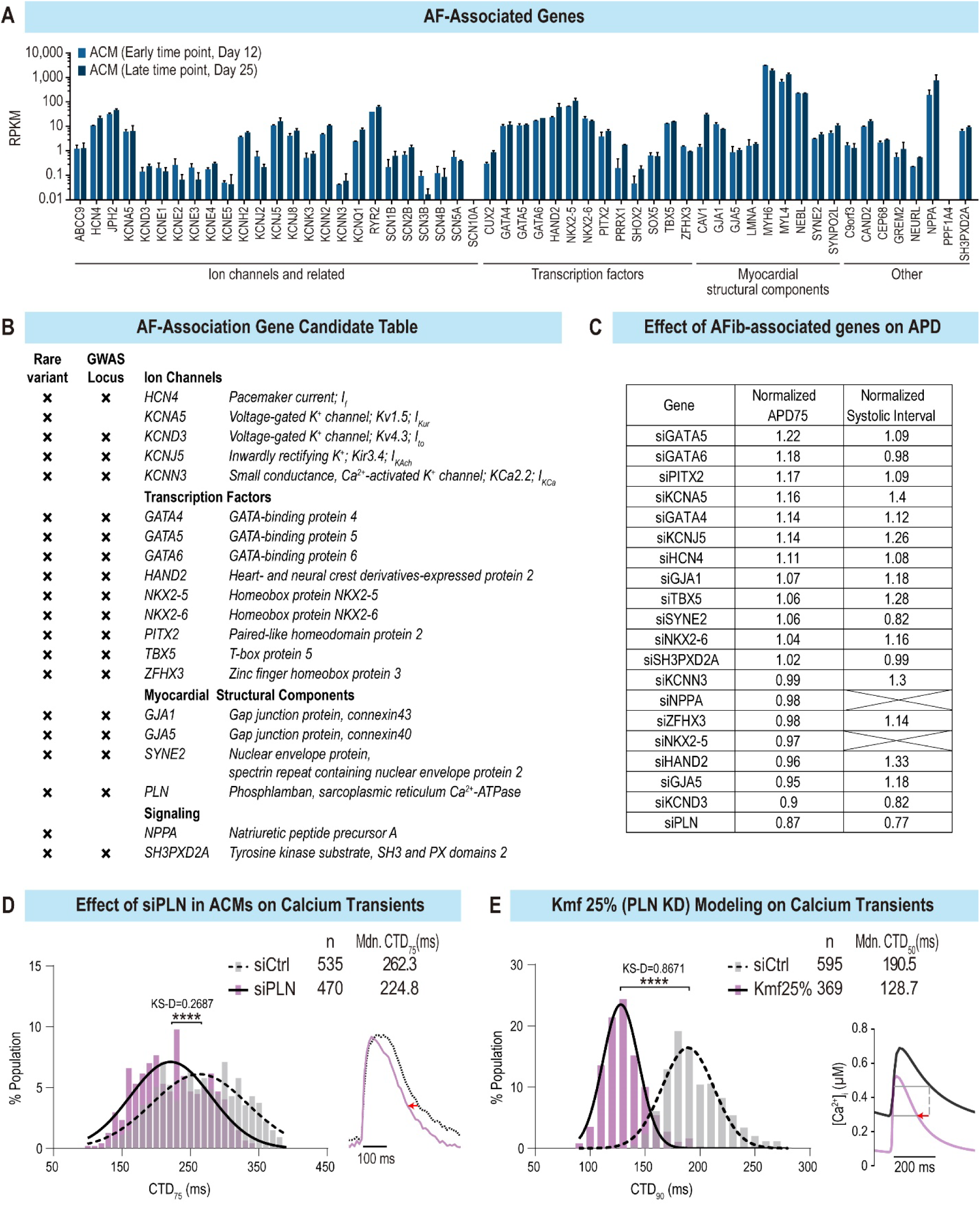
Selection of AF-associated genes for phenotypical characterization using our novel HT platform. **(A)** Histogram representing expression level (RPKM) of previously known AF-associated genes using RNAseq data from Day 12 and Day 25 ACMs. **(B)** Selection of 20 AF-genes harboring rare variant in familial AF studies and/or SNPs reported in GWAS studies. **(C)** Numerical values of data presented in heat map in Fig.3A. APD_75_ and systolic Intervals were normalized to their respective controls. In flies there is no homolog for NPPA and tinman/Nkx2.5 KD flies fail to develop hearts. **(D)** Histogram showing the distribution of calcium transient duration (CTD) values in siControl and siPLN condition in ACMs (right) and representative single calcium transients for both conditions (left). **(E)** Histogram of CTD generated from HAMs (left) and representative traces (right) in response toKmf25% condition (=PLN KD). P-value *** < 0.0001.

**Supplemental Figure 4.**
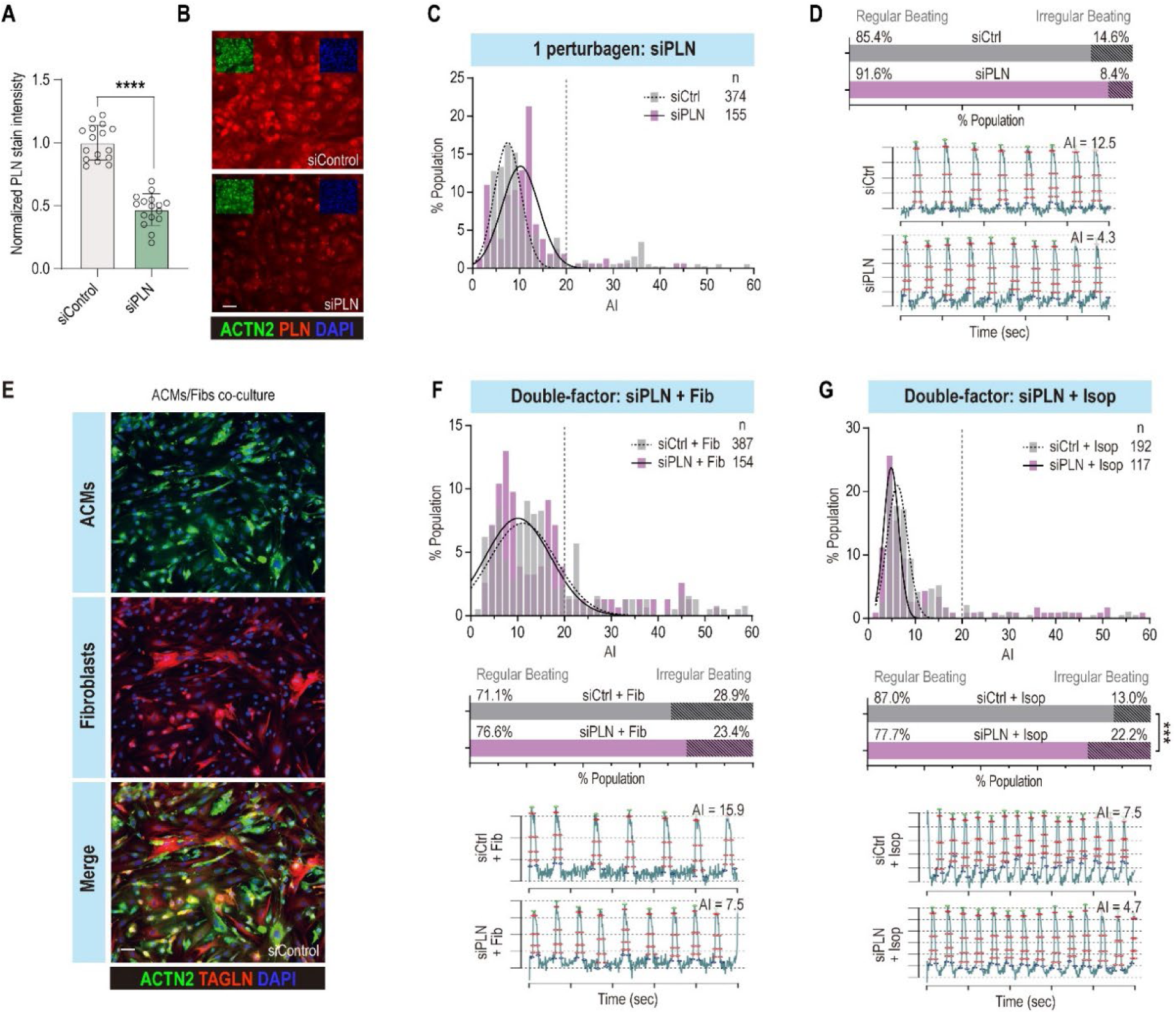
Testing of multiple perturbagens to trigger arrhythmia in ACMs. **(A)** Histogram showing the quantification of PLN staining intensity in siControl and siPLN conditions in ACMs. **(B)** Representative images of ACMs stained for actinin2 (ACTN2, green) to mark ACMs and for phospholamban (PLN, red) showing a significant reduction of PLN protein levels upon PLN KD. **(C)** Histogram showing the distribution of AI values from ACMs in siControl and siPLN conditions. **(D)** Quantification of irregular AP peak train (top). Representative AP traces in siControl and siPLN conditions. **(E)** Representative images of ACMs co-cultured with human dermal fibroblasts and stained with ACTN2 (cardiac, green) and TAGLN (fibroblasts, red). **(F)** Histogram showing the distribution of AI values from ACMs co-cultured with fibroblasts (Fib) in siControl and siPLN conditions (top). (Middle) Quantification of the percentage of ACMs ttwith irregular AP peak trains for each condition. (Bottom) Representative AP peak trains for each condition. **(G)** Histogram showing the distribution of AI values from ACMs treated with Isop in siControl and siPLN conditions (top). (Middle) Quantification of the percentage of ACMs with irregular AP peak trains for each condition. (Bottom) Representative AP peak trains for each condition.

**Supplemental Figure 5.**
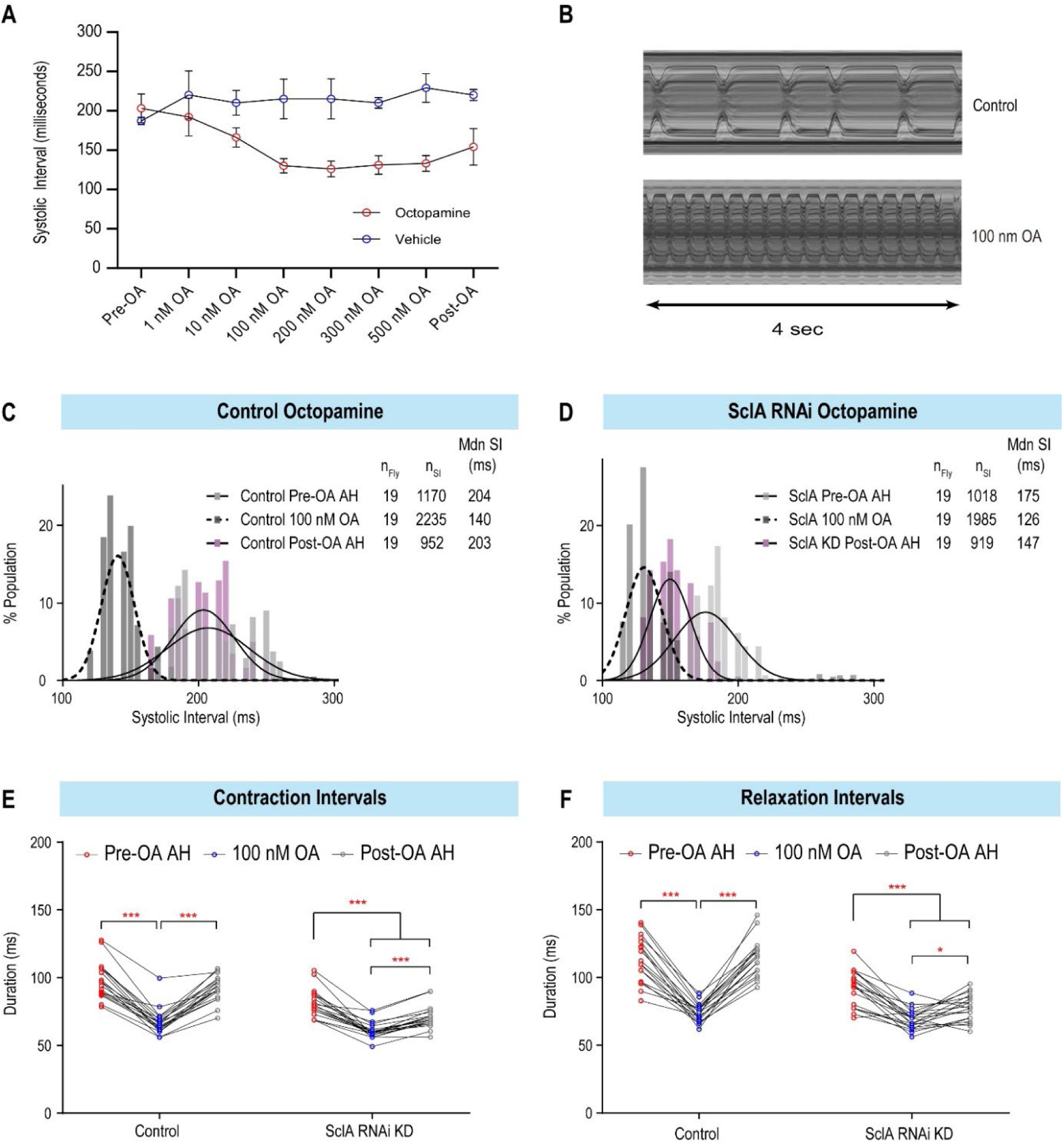
Optimization of octopamine treatment in flies. **(A)** Graph showing average SI values in response to escalating doses of OA in flies. **(B)** Representative M-modes from one heart before (top) and during application of 100nM OA (bottom) showing dramatic increases in heart rate. **(C)** Histograms showing the distribution of SI values in control hearts pre- OA pacing, during exposure to 100nM OA, and 15 min post-OA application. Distribution of SIs post-OA returns to that of pre-OA. **(D)** showing the distribution of SI values in SclA KD hearts pre- OA pacing, during exposure to 100nM OA, and 15 min post-OA application. Distribution of SIs post-OA are significantly shorter compared to pre-OA and are due to decreases in both **(E)** the contraction and **(F)** relaxation phases of the systolic intervals(p-value < 0.05, repeated measures 2-way ANOVA).

**Supplemental Fig. 6.**
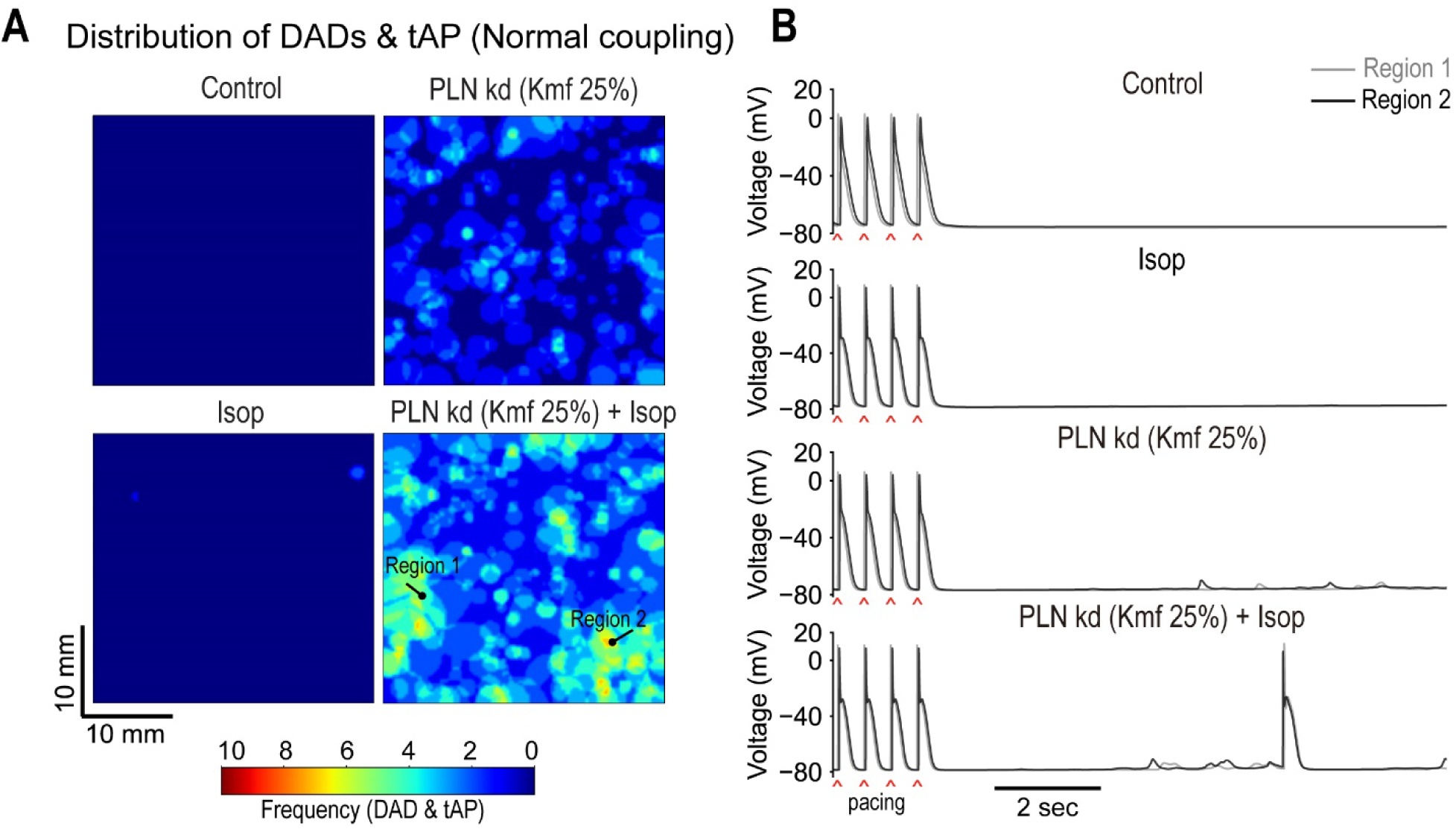
Effects of PLN kd (Kmf 25%) and Isop on the triggered activity in heterogeneous human atrial tissue with normal cell-to-cell coupling. Tissue was paced at 2 Hz for 10 s followed by a period of 10 s without stimulation. **(A)** Spatial distribution of DADs and tAPs in the atrial tissue for PLN KD (Kmf 25%) and after Isop treatment. **(B)** Superimposed traces of APs from two regions (marked in panel ***A***) of the atrial tissue with normal cell-to-cell coupling for each perturbation.

## References

Babu, G.J., Bhupathy, P., Timofeyev, V., Petrashevskaya, N.N., Reiser, P.J., Chiamvimonvat, N., and Periasamy, M. (2007). Ablation of sarcolipin enhances sarcoplasmic reticulum calcium transport and atrial contractility. Proceedings of the National Academy of Sciences of the United States of America 104, 17867–17872.

Birse, R.T., Choi, J., Reardon, K., Rodriguez, J., Graham, S., Diop, S., Ocorr, K., Bodmer, R., and Oldham, S. (2010). High-fat-diet-induced obesity and heart dysfunction are regulated by the TOR pathway in Drosophila. Cell Metab 12, 533–544.

Blice-Baum, A.C., Guida, M.C., Hartley, P.S., Adams, P.D., Bodmer, R., and Cammarato, A. (2019). As time flies by: Investigating cardiac aging in the short-lived Drosophila model. Biochim Biophys Acta Mol Basis Dis 1865, 1831–1844.

Bodmer, R. (1995). Heart development in Drosophila and its relationship to vertebrates. Trends in cardiovascular medicine 5, 21–28.

Brand, A.H., and Perrimon, N. (1993). Targeted gene expression as a means of altering cell fates and generating dominant phenotypes. Development 118, 401–415.

Burridge, P.W., Matsa, E., Shukla, P., Lin, Z.C., Churko, J.M., Ebert, A.D., Lan, F., Diecke, S., Huber, B., Mordwinkin, N.M., et al. (2014). Chemically defined generation of human cardiomyocytes. Nature methods 11, 855–860.

Cammarato, A., Ahrens, C.H., Alayari, N.N., Qeli, E., Rucker, J., Reedy, M.C., Zmasek, C.M., Gucek, M., Cole, R.N., Van Eyk, J.E., et al. (2011). A mighty small heart: the cardiac proteome of adult Drosophila melanogaster. PloS one 6, e18497.

Cammarato, A., Ocorr, S., and Ocorr, K. (2015). Enhanced assessment of contractile dynamics in Drosophila hearts. BioTechniques 58, 77–80.

Campuzano, O., and Brugada, R. (2009). Genetics of familial atrial fibrillation. Europace 11, 1267–1271.

Cantilina, T., Sagara, Y., Inesi, G., and Jones, L.R. (1993). Comparative studies of cardiac and skeletal sarcoplasmic reticulum ATPases. Effect of a phospholamban antibody on enzyme activation by Ca2+. The Journal of biological chemistry 268, 17018–17025.

Cerignoli, F., Charlot, D., Whittaker, R., Ingermanson, R., Gehalot, P., Savchenko, A., Gallacher, D.J., Towart, R., Price, J.H., McDonough, P.M., et al. (2012). High throughput measurement of Ca(2)(+) dynamics for drug risk assessment in human stem cell-derived cardiomyocytes by kinetic image cytometry. J Pharmacol Toxicol Methods 66, 246–256.

Chen, P.S., Chen, L.S., Fishbein, M.C., Lin, S.F., and Nattel, S. (2014). Role of the autonomic nervous system in atrial fibrillation: pathophysiology and therapy. Circulation research 114, 1500–1515.

Chen, Y.H., Xu, S.J., Bendahhou, S., Wang, X.L., Wang, Y., Xu, W.Y., Jin, H.W., Sun, H., Su, X.Y., Zhuang, Q.N., et al. (2003). KCNQ1 gain-of-function mutation in familial atrial fibrillation. Science 299, 251–254.

Cho, G.S., Lee, D.I., Tampakakis, E., Murphy, S., Andersen, P., Uosaki, H., Chelko, S., Chakir, K., Hong, I., Seo, K., et al. (2017). Neonatal Transplantation Confers Maturation of PSC-Derived Cardiomyocytes Conducive to Modeling Cardiomyopathy. Cell reports 18, 571–582.

Christophersen, I.E., Olesen, M.S., Liang, B., Andersen, M.N., Larsen, A.P., Nielsen, J.B., Haunso, S., Olesen, S.P., Tveit, A., Svendsen, J.H., et al. (2013). Genetic variation in KCNA5: impact on the atrial-specific potassium current IKur in patients with lone atrial fibrillation. Eur Heart J 34, 1517–1525.

Christophersen, I.E., Rienstra, M., Roselli, C., Yin, X., Geelhoed, B., Barnard, J., Lin, H., Arking, D.E., Smith, A.V., Albert, C.M., et al. (2017). Large-scale analyses of common and rare variants identify 12 new loci associated with atrial fibrillation. Nature genetics 49, 946–952.

Colman, M.A., Aslanidi, O.V., Kharche, S., Boyett, M.R., Garratt, C., Hancox, J.C., and Zhang, H. (2013). Pro-arrhythmogenic effects of atrial fibrillation-induced electrical remodelling: insights from the three-dimensional virtual human atria. J Physiol 591, 4249–4272.

Cripps, R.M., and Olson, E.N. (2002). Control of cardiac development by an evolutionarily conserved transcriptional network. Developmental biology 246, 14–28.

Cunningham, T.J., Yu, M.S., McKeithan, W.L., Spiering, S., Carrette, F., Huang, C.T., Bushway, P.J., Tierney, M., Albini, S., Giacca, M., et al. (2017). Id genes are essential for early heart formation. Genes & development.

Dal Molin, A., Baruzzo, G., and Di Camillo, B. (2017). Single-Cell RNA-Sequencing: Assessment of Differential Expression Analysis Methods. Front Genet 8, 62.

Delmans, M., and Hemberg, M. (2016). Discrete distributional differential expression (D3E)--a tool for gene expression analysis of single-cell RNA-seq data. BMC Bioinformatics 17, 110.

Denham, N.C., Pearman, C.M., Caldwell, J.L., Madders, G.W.P., Eisner, D.A., Trafford, A.W., and Dibb, K.M. (2018). Calcium in the Pathophysiology of Atrial Fibrillation and Heart Failure. Front Physiol 9, 1380.

Devalla, H.D., Gelinas, R., Aburawi, E.H., Beqqali, A., Goyette, P., Freund, C., Chaix, M.A., Tadros, R., Jiang, H., Le Bechec, A., et al. (2016). TECRL, a new life-threatening inherited arrhythmia gene associated with overlapping clinical features of both LQTS and CPVT. EMBO molecular medicine 8, 1390–1408.

Devalla, H.D., Schwach, V., Ford, J.W., Milnes, J.T., El-Haou, S., Jackson, C., Gkatzis, K., Elliott, D.A., Chuva de Sousa Lopes, S.M., Mummery, C.L., et al. (2015). Atrial-like cardiomyocytes from human pluripotent stem cells are a robust preclinical model for assessing atrial-selective pharmacology. EMBO molecular medicine 7, 394–410.

Diop, S.B., and Bodmer, R. (2015). Gaining Insights into Diabetic Cardiomyopathy from Drosophila. Trends Endocrinol Metab 26, 618–627.

Du, X., Dong, J., and Ma, C. (2017). Is Atrial Fibrillation a Preventable Disease? Journal of the American College of Cardiology 69, 1968–1982.

Dzeshka, M.S., Lip, G.Y., Snezhitskiy, V., and Shantsila, E. (2015). Cardiac Fibrosis in Patients With Atrial Fibrillation: Mechanisms and Clinical Implications. Journal of the American College of Cardiology 66, 943–959.

Elmen, L., Volpato, C.B., Kervadec, A., Pineda, S., Kalvakuri, S., Alayari, N.N., Foco, L., Pramstaller, P.P., Ocorr, K., Rossini, A., et al. (2020). Silencing of CCR4-NOT complex subunits affect heart structure and function. Dis Model Mech.

Fatkin, D., Santiago, C.F., Huttner, I.G., Lubitz, S.A., and Ellinor, P.T. (2017). Genetics of Atrial Fibrillation: State of the Art in 2017. Heart Lung Circ 26, 894–901.

Feng, Y., Mitchison, T.J., Bender, A., Young, D.W., and Tallarico, J.A. (2009). Multi-parameter phenotypic profiling: using cellular effects to characterize small-molecule compounds. Nat Rev Drug Discov 8, 567–578.

Fink, M., Callol-Massot, C., Chu, A., Ruiz-Lozano, P., Izpisua Belmonte, J.C., Giles, W., Bodmer, R., and Ocorr, K. (2009). A new method for detection and quantification of heartbeat parameters in Drosophila, zebrafish, and embryonic mouse hearts. BioTechniques 46, 101–113.

Gaber, N., Gagliardi, M., Patel, P., Kinnear, C., Zhang, C., Chitayat, D., Shannon, P., Jaeggi, E., Tabori, U., Keller, G., et al. (2013). Fetal reprogramming and senescence in hypoplastic left heart syndrome and in human pluripotent stem cells during cardiac differentiation. Am J Pathol 183, 720–734.

Geng, M., Lin, A., and Nguyen, T.P. (2020). Revisiting Antiarrhythmic Drug Therapy for Atrial Fibrillation: Reviewing Lessons Learned and Redefining Therapeutic Paradigms. Front Pharmacol 11, 581837.

Grandi, E., Pandit, S.V., Voigt, N., Workman, A.J., Dobrev, D., Jalife, J., and Bers, D.M. (2011). Human atrial action potential and Ca2+ model: sinus rhythm and chronic atrial fibrillation. Circulation research 109, 1055–1066.

Hodgson-Zingman, D.M., Karst, M.L., Zingman, L.V., Heublein, D.M., Darbar, D., Herron, K.J., Ballew, J.D., de Andrade, M., Burnett, J.C., Jr., and Olson, T.M. (2008). Atrial natriuretic peptide frameshift mutation in familial atrial fibrillation. N Engl J Med 359, 158–165.

Jaiswal, A., and Goldbarg, S. (2014). Dofetilide induced torsade de pointes: mechanism, risk factors and management strategies. Indian Heart J 66, 640–648.

Johnson, E., Ringo, J., and Dowse, H. (1997). Modulation of Drosophila heartbeat by neurotransmitters. J Comp Physiol B 167, 89–97.

Kaese, S., and Verheule, S. (2012). Cardiac electrophysiology in mice: a matter of size. Front Physiol 3, 345.

Klassen, M.P., Peters, C.J., Zhou, S., Williams, H.H., Jan, L.Y., and Jan, Y.N. (2017). Age-dependent diastolic heart failure in an in vivo Drosophila model. eLife 6.

Lip, G.Y., Fauchier, L., Freedman, S.B., Van Gelder, I., Natale, A., Gianni, C., Nattel, S., Potpara, T., Rienstra, M., Tse, H.F., et al. (2016). Atrial fibrillation. Nat Rev Dis Primers 2, 16016.

Liu, P., and Miller, E.W. (2020). Electrophysiology, Unplugged: Imaging Membrane Potential with Fluorescent Indicators. Acc Chem Res 53, 11–19.

Lozano-Velasco, E., Hernandez-Torres, F., Daimi, H., Serra, S.A., Herraiz, A., Hove-Madsen, L., Aranega, A., and Franco, D. (2016). Pitx2 impairs calcium handling in a dose-dependent manner by modulating Wnt signalling. Cardiovascular research 109, 55–66.

Marczenke, M., Fell, J., Piccini, I., Ropke, A., Seebohm, G., and Greber, B. (2017a). Generation and cardiac subtype-specific differentiation of PITX2-deficient human iPS cell lines for exploring familial atrial fibrillation. Stem Cell Res 21, 26–28.

Marczenke, M., Piccini, I., Mengarelli, I., Fell, J., Ropke, A., Seebohm, G., Verkerk, A.O., and Greber, B. (2017b). Cardiac Subtype-Specific Modeling of Kv1.5 Ion Channel Deficiency Using Human Pluripotent Stem Cells. Front Physiol 8, 469.

McKeithan, W.L., Savchenko, A., Yu, M.S., Cerignoli, F., Bruyneel, A.A.N., Price, J.H., Colas, A.R., Miller, E.W., Cashman, J.R., and Mercola, M. (2017). An Automated Platform for Assessment of Congenital and Drug-Induced Arrhythmia with hiPSC-Derived Cardiomyocytes. Front Physiol 8, 766.

Mohr, S.E., Hu, Y., Kim, K., Housden, B.E., and Perrimon, N. (2014). Resources for functional genomics studies in Drosophila melanogaster. Genetics 197, 1–18.

Morotti, S., and Grandi, E. (2017). Logistic regression analysis of populations of electrophysiological models to assess proarrythmic risk. MethodsX 4, 25–34.

Morotti, S., Nieves-Cintron, M., Nystoriak, M.A., Navedo, M.F., and Grandi, E. (2017). Predominant contribution of L-type Cav1.2 channel stimulation to impaired intracellular calcium and cerebral artery vasoconstriction in diabetic hyperglycemia. Channels (Austin) 11, 340–346.

Murphy, S.A., Miyamoto, M., Kervadec, A., Kannan, S., Tampakakis, E., Kambhampati, S., Lin, B.L., Paek, S., Andersen, P., Lee, D.I., et al. (2021). PGC1/PPAR drive cardiomyocyte maturation at single cell level via YAP1 and SF3B2. Nature communications 12, 1648.

Nadadur, R.D., Broman, M.T., Boukens, B., Mazurek, S.R., Yang, X., van den Boogaard, M., Bekeny, J., Gadek, M., Ward, T., Zhang, M., et al. (2016). Pitx2 modulates a Tbx5-dependent gene regulatory network to maintain atrial rhythm. Sci Transl Med 8, 354ra115.

Nassal, M.M., Wan, X., Laurita, K.R., and Cutler, M.J. (2015). Atrial SERCA2a Overexpression Has No Affect on Cardiac Alternans but Promotes Arrhythmogenic SR Ca2+ Triggers. PloS one 10, e0137359.

Neely, G.G., Kuba, K., Cammarato, A., Isobe, K., Amann, S., Zhang, L., Murata, M., Elmen, L., Gupta, V., Arora, S., et al. (2010). A global in vivo Drosophila RNAi screen identifies NOT3 as a conserved regulator of heart function. Cell 141, 142–153.

Ng, S.Y., Wong, C.K., and Tsang, S.Y. (2010). Differential gene expressions in atrial and ventricular myocytes: insights into the road of applying embryonic stem cell-derived cardiomyocytes for future therapies. Am J Physiol Cell Physiol 299, C1234–1249.

Ni, H., Morotti, S., and Grandi, E. (2018). A Heart for Diversity: Simulating Variability in Cardiac Arrhythmia Research. Front Physiol 9, 958.

Ni, H., Whittaker, D.G., Wang, W., Giles, W.R., Narayan, S.M., and Zhang, H. (2017). Synergistic Anti-arrhythmic Effects in Human Atria with Combined Use of Sodium Blockers and Acacetin. Front Physiol 8, 946.

Nielsen, J.B., Fritsche, L.G., Zhou, W., Teslovich, T.M., Holmen, O.L., Gustafsson, S., Gabrielsen, M.E., Schmidt, E.M., Beaumont, R., Wolford, B.N., et al. (2018a). Genome-wide Study of Atrial Fibrillation Identifies Seven Risk Loci and Highlights Biological Pathways and Regulatory Elements Involved in Cardiac Development. Am J Hum Genet 102, 103–115.

Nielsen, J.B., Graff, C., Pietersen, A., Lind, B., Struijk, J.J., Olesen, M.S., Haunso, S., Gerds, T.A., Svendsen, J.H., Kober, L., et al. (2013). J-shaped association between QTc interval duration and the risk of atrial fibrillation: results from the Copenhagen ECG study. Journal of the American College of Cardiology 61, 2557–2564.

Nielsen, J.B., Thorolfsdottir, R.B., Fritsche, L.G., Zhou, W., Skov, M.W., Graham, S.E., Herron, T.J., McCarthy, S., Schmidt, E.M., Sveinbjornsson, G., et al. (2018b). Biobank-driven genomic discovery yields new insight into atrial fibrillation biology. Nature genetics 50, 1234–1239.

Nielsen, T., Kervadec, A., Missinato, M.A., Lynott, M., Zeng, X.-X.I., Berenguer, M., Walls, S.M., Schroeder, A., Birker, K., Duester, G., et al. (2022). Functional analysis across model systems implicates ribosomal proteins in growth and proliferation defects associated with hypoplastic left heart syndrome. medRxiv, 2022.2007.2001.22277112.

Nishimura, M., Ocorr, K., Bodmer, R., and Cartry, J. (2011). Drosophila as a model to study cardiac aging. Exp Gerontol 46, 326–330.

Ocorr, K., Fink, M., Cammarato, A., Bernstein, S., and Bodmer, R. (2009). Semi-automated Optical Heartbeat Analysis of small hearts. Journal of visualized experiments : JoVE.

Ocorr, K., Perrin, L., Lim, H.Y., Qian, L., Wu, X., and Bodmer, R. (2007a). Genetic control of heart function and aging in Drosophila. Trends in cardiovascular medicine 17, 177–182.

Ocorr, K., Reeves, N.L., Wessells, R.J., Fink, M., Chen, H.S., Akasaka, T., Yasuda, S., Metzger, J.M., Giles, W., Posakony, J.W., et al. (2007b). KCNQ potassium channel mutations cause cardiac arrhythmias in Drosophila that mimic the effects of aging. Proceedings of the National Academy of Sciences of the United States of America 104, 3943–3948.

Ocorr, K., Vogler, G., and Bodmer, R. (2014). Methods to assess Drosophila heart development, function and aging. Methods 68, 265–272.

Ocorr, K., Zambon, A., Nudell, Y., Pineda, S., Diop, S., Tang, M., Akasaka, T., and Taylor, E. (2017). Age-dependent electrical and morphological remodeling of the Drosophila heart caused by hERG/seizure mutations. PLoS genetics 13, e1006786.

Olson, T.M., Alekseev, A.E., Liu, X.K., Park, S., Zingman, L.V., Bienengraeber, M., Sattiraju, S., Ballew, J.D., Jahangir, A., and Terzic, A. (2006). Kv1.5 channelopathy due to KCNA5 loss-of-function mutation causes human atrial fibrillation. Hum Mol Genet 15, 2185–2191.

Paredes, R.M., Etzler, J.C., Watts, L.T., Zheng, W., and Lechleiter, J.D. (2008). Chemical calcium indicators. Methods 46, 143–151.

Periasamy, M., Bhupathy, P., and Babu, G.J. (2008). Regulation of sarcoplasmic reticulum Ca2+ ATPase pump expression and its relevance to cardiac muscle physiology and pathology. Cardiovascular research 77, 265–273.

Roselli, C., Chaffin, M.D., Weng, L.C., Aeschbacher, S., Ahlberg, G., Albert, C.M., Almgren, P., Alonso, A., Anderson, C.D., Aragam, K.G., et al. (2018). Multi-ethnic genome-wide association study for atrial fibrillation. Nature genetics.

Roselli, C., Rienstra, M., and Ellinor, P.T. (2020). Genetics of Atrial Fibrillation in 2020: GWAS, Genome Sequencing, Polygenic Risk, and Beyond. Circulation research 127, 21–33.

Salick, M.R., Napiwocki, B.N., Sha, J., Knight, G.T., Chindhy, S.A., Kamp, T.J., Ashton, R.S., and Crone, W.C. (2014). Micropattern width dependent sarcomere development in human ESC-derived cardiomyocytes. Biomaterials 35, 4454–4464.

Schuttler, D., Bapat, A., Kaab, S., Lee, K., Tomsits, P., Clauss, S., and Hucker, W.J. (2020). Animal Models of Atrial Fibrillation. Circulation research 127, 91–110.

Sellin, J., Albrecht, S., Kolsch, V., and Paululat, A. (2006). Dynamics of heart differentiation, visualized utilizing heart enhancer elements of the Drosophila melanogaster bHLH transcription factor Hand. Gene Expr Patterns 6, 360–375.

Simmerman, H.K., and Jones, L.R. (1998). Phospholamban: protein structure, mechanism of action, and role in cardiac function. Physiol Rev 78, 921–947.

Sobie, E.A. (2009). Parameter sensitivity analysis in electrophysiological models using multivariable regression. Biophys J 96, 1264–1274.

Sujkowski, A., Ramesh, D., Brockmann, A., and Wessells, R. (2017). Octopamine Drives Endurance Exercise Adaptations in Drosophila. Cell reports 21, 1809–1823.

Takahashi, K., Tanabe, K., Ohnuki, M., Narita, M., Ichisaka, T., Tomoda, K., and Yamanaka, S. (2007). Induction of pluripotent stem cells from adult human fibroblasts by defined factors. Cell 131, 861–872.

Takahashi, K., and Yamanaka, S. (2006). Induction of pluripotent stem cells from mouse embryonic and adult fibroblast cultures by defined factors. Cell 126, 663–676.

Teh, A.W., Kistler, P.M., Lee, G., Medi, C., Heck, P.M., Spence, S.J., Sparks, P.B., Morton, J.B., and Kalman, J.M. (2012). Electroanatomic remodeling of the left atrium in paroxysmal and persistent atrial fibrillation patients without structural heart disease. J Cardiovasc Electrophysiol 23, 232–238.

Temple, J., Frias, P., Rottman, J., Yang, T., Wu, Y., Verheijck, E.E., Zhang, W., Siprachanh, C., Kanki, H., Atkinson, J.B., et al. (2005). Atrial fibrillation in KCNE1-null mice. Circulation research 97, 62–69.

Uhlen, M., Fagerberg, L., Hallstrom, B.M., Lindskog, C., Oksvold, P., Mardinoglu, A., Sivertsson, A., Kampf, C., Sjostedt, E., Asplund, A., et al. (2015). Proteomics. Tissue-based map of the human proteome. Science 347, 1260419.

Uosaki, H., Cahan, P., Lee, D.I., Wang, S., Miyamoto, M., Fernandez, L., Kass, D.A., and Kwon, C. (2015). Transcriptional Landscape of Cardiomyocyte Maturation. Cell reports 13, 1705–1716.

van Ouwerkerk, A.F., Bosada, F.M., van Duijvenboden, K., Hill, M.C., Montefiori, L.E., Scholman, K.T., Liu, J., de Vries, A.A.F., Boukens, B.J., Ellinor, P.T., et al. (2019). Identification of atrial fibrillation associated genes and functional non-coding variants. Nature communications 10, 4755.

Vogler, G., and Ocorr, K. (2009). Visualizing the beating heart in Drosophila. Journal of visualized experiments : JoVE.

Voigt, N., Heijman, J., Wang, Q., Chiang, D.Y., Li, N., Karck, M., Wehrens, X.H.T., Nattel, S., and Dobrev, D. (2014). Cellular and molecular mechanisms of atrial arrhythmogenesis in patients with paroxysmal atrial fibrillation. Circulation 129, 145–156.

Wakili, R., Voigt, N., Kaab, S., Dobrev, D., and Nattel, S. (2011). Recent advances in the molecular pathophysiology of atrial fibrillation. The Journal of clinical investigation 121, 2955–2968.

Wang, J., Klysik, E., Sood, S., Johnson, R.L., Wehrens, X.H., and Martin, J.F. (2010). Pitx2 prevents susceptibility to atrial arrhythmias by inhibiting left-sided pacemaker specification. Proceedings of the National Academy of Sciences of the United States of America 107, 9753–9758.

Weng, L.C., Choi, S.H., Klarin, D., Smith, J.G., Loh, P.R., Chaffin, M., Roselli, C., Hulme, O.L., Lunetta, K.L., Dupuis, J., et al. (2017). Heritability of Atrial Fibrillation. Circ Cardiovasc Genet 10.

Workman, A.J. (2010). Cardiac adrenergic control and atrial fibrillation. Naunyn Schmiedebergs Arch Pharmacol 381, 235–249.

Xintarakou, A., Tzeis, S., Psarras, S., Asvestas, D., and Vardas, P. (2020). Atrial fibrosis as a dominant factor for the development of atrial fibrillation: facts and gaps. Europace 22, 342–351.

Yang, X., Pabon, L., and Murry, C.E. (2014). Engineering adolescence: maturation of human pluripotent stem cell-derived cardiomyocytes. Circulation research 114, 511–523.

Yu, M.S., Spiering, S., and Colas, A.R. (2018). Generation of First Heart Field-like Cardiac Progenitors and Ventricular-like Cardiomyocytes from Human Pluripotent Stem Cells. Journal of visualized experiments : JoVE.

Zhang, M., Hill, M.C., Kadow, Z.A., Suh, J.H., Tucker, N.R., Hall, A.W., Tran, T.T., Swinton, P.S., Leach, J.P., Margulies, K.B., et al. (2019). Long-range Pitx2c enhancer-promoter interactions prevent predisposition to atrial fibrillation. Proceedings of the National Academy of Sciences of the United States of America 116, 22692–22698.

Zhao, D., Lei, W., and Hu, S. (2021). Cardiac organoid - a promising perspective of preclinical model. Stem Cell Res Ther 12, 272.

