## Supplementary material for "Multiplatform Modeling of Atrial Fibrillation Identifies Phospholamban as Central Regulator of Cardiac Rhythm": All supplemental Figures

**Supplemental Table 1**

| CATEGORY | GENE | FLY ORTHOLOG | Reference | FlyAtlas2 |
|---|---|---|---|---|
| CHANNEL | **ABCC9** | **dSur** | Akasaka, et al 2006 -PMID: 16882722; Eleftherianos et al, 2011 - PMID: 21719711 | **-** |
| CHANNEL | **HCN4** | **Ih** | Monier, et al, 2005 - PMID 16284119 | **+** |
| CHANNEL | **JPH2** | **junctophilin** | | **+** |
| CHANNEL | **KCNA5** | **Shaker** | Ocorr et al, 2017 - PMID: 28542428 | **-** |
| CHANNEL | **KCND3** | **Shal** | Ocorr et al, 2017 - PMID: 28542428 | **-** |
| CHANNEL | KCNE1 - 5 | No ortholog | KCNQ works without MinK | |
| CHANNEL | **KCNH2** | **seizure** | Ocorr et al, 2017 - PMID:28542428 | **-** |
| CHANNEL | **KCNJ2, 5, 8** | **Irk** | Ocorr et al, 2017 - PMID:28542428 | **-** |
| CHANNEL | **KCNK3** | **ork / sandman** | LaLevee et al – PMID: 16890525<br>Klassen et al - PMID: 28328397 | **- / -** |
| CHANNEL | **KCNN2, 3** | **SK** | | **+** |
| CHANNEL | **KCNMA** | **BK – not tested in ACMs?** | Pineda et al, - PMID: 33629867 | **+** |
| CHANNEL | **KCNQ1** | **KCNQ** | Ocorr et al, 2007 - PMID: 17360457 | **-** |
| CHANNEL | **RYR2** | **RyR** | Lin et al, PMID: 21493892 | **+** |
| CHANNEL | **SCN1 - 5** | **nap** | Dowse et al, PMID: 8719771<br>Ganetsky, PMID: 2420953 | **+** |
| TF | **CUX2** | **cut** | Blochlinger et al, PMID 8330519<br>Zappia et al, PMID 32815271 | **-** |
| TF | **GATA4 / 5 / 6** | **pnr / grn / GATAd** | Klinedinst & Bodmer, 2003 - PMID: 12756184 | **+ / + / +** |
| TF | **HAND2** | **Hand** | Kolsch and Paululat, 2002 - PMID: 12424518 Jonhson et al 2011 - PMID: 21965617 | **+** |
| TF | **NKX2-5 / 2-6** | **tin** | Bodmer et al 1990 - PMID: 7915669 | **-** |
| TF | PITX2 | Ptx1 | | **-** |
| TF | PRRX1 | CG9876 | | **-** |
| TF | SHOX2 | CG34367 | | **-** |
| TF | **SOX5** | **Sox102F** | | **+** |
| TF | **TBX5** | **Bifid / Doc1 / Doc2 / Doc3** | Bi - Ahmad et al 2012 - PMID: 22814603<br>DOC -Reim et al 2003 - PMID: 12783790 Berkeley Drosophila Genome Project | **- / - /- /-** |
| TF | ZFHX3 | zfh2 | | **-** |
| MYOCARDIAL | CAV1 | No ortholog | | |
| MYOCARDIAL | **GJA1** | **CG11459 / 26-29-p / CG4847** | Cammarato et al 2011 - PMID: 21541028 | **+** |
| MYOCARDIAL | GJA5 | No ortholog | | |

| | | | | |
|---|---|---|---|---|
| MYOCARDIAL | **LMNA** | **LamC / Lam** | Cammarato et al 2011 - PMID: 21541028 | **+** |
| MYOCARDIAL | **MYH6** | **Mhc** | Lovato et al, 2002 - PMID: 12397110 Cammarato et al, 2011 - PMID: 21541028 | **+** |
| MYOCARDIAL | **MYL4** | **Mlc-c / Mlc1** | Cammarato et al 2011 - PMID: 21541028 | **+** |
| MYOCARDIAL | **NEBL** | **Lasp** | Cammarato et al 2011 - PMID: 21541028 | **+** |
| MYOCARDIAL | **SYNE2** | **Msp300** | Cammarato et al 2011 - PMID: 21541028 | **+** |
| MYOCARDIAL | **SYNPO2L** | **CG1674** | Cammarato et al 2011 - PMID: 21541028 | **+** |
| OTHER | **C9ORF3** | **CG10602** | | **+** |
| OTHER | **CAND2** | **Cand1** | | **+** |
| OTHER | CEP68 | No ortholog | | |
| OTHER | GREM2 | No ortholog | | |
| OTHER | **NEURL** | **neur** | | **+** |
| OTHER | NPPA | No ortholog | | |
| OTHER | **SH3PXD2A** | **cindr / Nipped-A** | Cammarato et al 2011 - PMID: 21541028 | **+ / +** |

**Supplemental Table 1 –** Human AF candidate genes tested in ACMs (Supplemental Fig. 2) are shown with their *Drosophila* orthologs. Studies demonstrating the presence and/or function of these orthologs in the fly heart are listed and genes common to both ACMs and fly hearts are bolded. Data from tissue-specific RNA Seq analysis and curated in FlyAtlas2 (https://flyatlas.gla.ac.uk/FlyAtlas2/) shows cardiac expression for approximately half of the genes in the table (indicated by +). Note that many of the ion channels and transcription factors (TF) that have been shown to be functional in the heart did not show up in the Fly Atlas dataset and none showed up in the cardiac proteomic analysis by Cammarato et al (2011, PMID: 21541028), likely because expression in the heart is too low relative to other structural genes (e.g. Myosin heavy chain, Mhc).

**Supplemental Table 2**

| Fly RNAi | Human Ortholog | Stock # | Stock #2 | Source |
|---|---|---|---|---|
| **Bifid** | TBX 2,3 | 100598 | 330228 | VDRC |
| **Cindr** | SH3KBP1 | 38854 | 330411 | VDRC |
| **Doc1** | TBX6 | 16746 | 104927 | VDRC |
| **Doc2** | TBX6 | 103431 | | VDRC |
| **Doc3** | TBX6 | 30550 | 104922 | VDRC |
| **GATAd** | GATA1 | 100389 | | VDRC |
| **GATAe** | GATA4 | 10418 | | VDRC |
| **Grain** | GATA2,3 | 105192 | 330376 | VDRC |
| **Hand** | Hand | 23306 | 330058 | VDRC |
| **Ih** | HCN2-4 | 110274 | 29574 | VDRC |
| **Irk1** | KCNJ2,4,12,18 | 107389 | 28431 | VDRC |
| **Irk2** | KCNJ2,4,12,18 | 4341 | 108140 | VDRC |
| **Irk3** | KCNJ10,15 | 3886 | 101174 | VDRC |
| **MSP300** | SYNE1 | 107183 | 25906 | VDRC |
| **Pnr** | GATA4,5,6 | 6224 | 101522 | VDRC |
| **Pnr** | GATA4,5,6 | 34659 | 33744 | BDSC |
| **Ptx1** | Ptx1-3 | 19831 | 107785 | VDRC |
| **SclA** | PLN | 28957 | | BDSC |
| **SclA/B** | PLN | 62935 | 28957 | BDSC |
| **Scro** | NKX2.1,2.4 | 33902 | 330398 | VDRC |
| **Sh** | KCNA1-5 | 23673 | 104474 | VDRC |
| **Shal** | KCND1-3 | 103363 | 330383 | VDRC |
| **Sk** | KCNN1-3 | 2855 | 7052 | VDRC |
| **Sk** | KCNN1-3 | 27238 | 53881 | BDSC |
| **Tin** | NKX2.5 | 190512 | 101825 | VDRC |

| | | | | |
|---|---|---|---|---|
| **Twist** | TWIST1, 2 | 37091 | 37092 | VDRC |
| **Zfh-2** | ZFH2-4 | 13305 | 110784 | VDRC |

**Supplemental Table 2 –** Human AF candidate genes tested in the fly heart (Supplemental Fig. 2) are listed with their human orthologs and the stock center ID number.

**Supplemental Table 3**

| Parameter | Note |
|---|---|
| $G_{Na}$ | Fast $Na^+$ current, maximal conductance |
| $G_{CaL}$ | L-type $Ca^{2+}$current, maximal conductance |
| $G_{to}$ | Transient outward $K^+$ current, maximal conductance |
| $G_{Kur}$ | Ultra-rapid delayed rectifier $K^+$ current, maximal conductance |
| $G_{Kr}$ | Rapid delayed rectifier $K^+$ current, maximal conductance |
| $G_{Ks}$ | Slow delayed rectifier $K^+$ current, maximal conductance |
| $G_{K1}$ | Inward rectifier $K^+$ current, maximal conductance |
| $G_{Kp}$ | Conductance of the plateau $K^+$ current |
| $G_{NaB}$ | Background $Na^+$ current, maximal conductance |
| $G_{CaB}$ | Background $Ca^{2+}$ current, maximal conductance |
| $G_{CaP}$ | Sarcoplasmic $Ca^{2+}$ pump current, maximal pump rate |
| $G_{ClCa}$ | $Ca^{2+}$ activated $Cl^-$ current, maximal conductance |
| $G_{ClB}$ | Background $Cl^-$ current, maximal conductance |
| $V_{NCX}$ | $Na^+/Ca^{2+}$ exchange current, maximal exchange rate |
| $V_{NaK}$ | $Na^+/K^+$ pump current, maximal pump rate |
| $V_{SERCA}$ | Rate of the SERCA pump |
| $V_{RyR,Rel}$ | Rate of the SR $Ca^{2+}$ release via ryanodine receptors |
| $V_{RyR,Leak}$ | Rate of the SR $Ca^{2+}$ leak via ryanodine receptors |

**Supplemental Table 3**. Glossary for model parameters that were perturbed for constructing populations of human atrial models.
